# A comparative analysis of planarian regeneration specificity reveals tissue polarity contributions of the axial cWnt signalling gradient

**DOI:** 10.1101/2024.12.01.626254

**Authors:** James P. Cleland, Hanh T.-K. Vu, Johanna E. M. Dickmann, Andrei Rozanski, Steffen Werner, Andrea Schuhmann, Anna Shevchenko, Jochen C. Rink

## Abstract

Planarians exhibit remarkable whole-body regeneration abilities. The formation of heads at forward-facing wounds and tails at rearward-facing wounds suggests an intrinsic tissue polarity guiding regeneration. While the underlying mechanisms remain unclear, reports of double-headed regenerates from increasingly narrow tissue fragments have long been hypothesised to reflect gradient-based polarity specification. Here, we systematically re-examine this hypothesis in the modern model species *Schmidtea mediterranea* and a representative of the genus likely used in the original studies, *Girardia sinensis*. While we never observed double-heads in *S. mediterranea*, *G. sinensis* readily regenerated double-heads in a manner dependent on piece length, anatomical position and body size. We found that the reduced regeneration robustness of *G. sinensis* was partially explained by wound site-symmetric expression of the head determinant *notum*, which is highly anterior-specific in *S. mediterranea*. Exploring what else might mediate head/tail regeneration specificity in *G. sinensis*, we examined the role of the conserved tail-to-head cWnt signalling gradient. By developing a time-resolved pharmacological approach to reduce the cWnt gradient slope without affecting wound-induced cWnt signalling dynamics, we observed an increased incidence of double-headed regenerates. In addition, the body size-dependence of double-head formation correlated with the decreasing steepness of the cWnt gradient due to scaling. Taken together, our results indicate that the slope of the cWnt gradient may contribute to planarian head/tail regeneration specificity. Furthermore, they suggest that planarian tissue polarity is composed of multiple parallely-acting polarity cues, the differential reliance on which contributes to the observed interspecies variation in regeneration specificity.

## Introduction

The establishment and maintenance of species-specific shape and proportions remains a fascinating challenge in biology. In most animals, the body plan emerges during the course of embryonic development via a hierarchical cascade of developmental processes that is set in motion by symmetry-breaking events at the earliest stages of development. However, adults of many animal groups, including hydra, planaria, salamanders and fishes, can also regrow lost or injured body parts through the process of regeneration^1^. In sharp contrast to the fertilised zygote as invariant starting point of embryonic development, regeneration initiates from the unpredictable remnants of injury to restore precisely the missing body part and no more or less. For instance, when the head and the tail of a planarian are amputated, the remaining trunk piece regenerates one head and one tail to once again complete the species-specific body plan^2–4^. Despite progress in recent years in multiple model systems^5^, the mechanistic basis of “regeneration specificity” as the cellular and molecular mechanisms that so robustly ensure the re-completion of the species-specific body plan irrespective of the nature of the injury, remain incompletely understood in any system.

The ability of many planarian species to rapidly regenerate their entire body almost irrespective of the anatomical origin, shape or size of the regenerating tissue piece^2–4^ makes them a powerful model system for addressing the mechanistic underpinnings of regeneration specificity. Particularly the question of how regenerating pieces succeed in robustly regenerating both head and tail has been subject to intense investigation for more than a century. The fact that the new head and tail regenerate in alignment with the original body axes (i.e., heads from forward- and tails from rearward-facing wounds) indicates that an intrinsic tissue polarisation is a core aspect of planarian regeneration specificity. Interestingly, T.H. Morgan noted already in 1898 the breakdown of regeneration specificity in very narrow cross-pieces (e.g., transverse amputations), which he reported to often form bipolar animals with heads regenerating from both wounds^6,7^. Attempting to rationalise the apparent piece-length dependence of double-head formation, he proposed an underlying “gradation of materials” along the anteroposterior (A/P) axis as mechanistic polarity manifestation. Specifically, he envisaged the resulting difference in “material” between the anterior and posterior wounds of a crosspiece as instructive for head/tail regeneration, and hence the breakdown of regeneration specificity in very narrow crosspieces once this difference becomes too small^8^. Further observations broadly consistent with the gradient hypothesis include the systematic decrease of head regeneration rates towards the posterior in many species^8–10^ and the complete head regeneration failures of multiple planarian species in the posterior body half^9,11–14^. Importantly, the existence of organism-scale gene expression and signalling pathway activity gradients along the planarian A/P axis has meanwhile been demonstrated. These may include a head-to-tail gradient of extracellular-signal regulated kinase (Erk) activity^13,15^ and a demonstrated tail-to-head gradient of canonical Wnt (cWnt) signalling^16,17^. Moreover, high cWnt signalling is necessary and sufficient for tail specification, while cWnt inhibition is necessary and sufficient for head specification^18–20^. During regeneration, the selective expression of the cWnt antagonist *notum* at anterior-facing wounds is indispensable for head regeneration^21,22^. Although some of the signalling pathways contributing to the anterior specificity of *notum* expression have meanwhile been identified (including cWnt signalling itself)^21,23,24^ and *notum* expression has been localised to longitudinal muscle fibres^21,22^, the nature of the head/tail specifying polarity cue and whether or not it involves signalling gradients remains fundamentally unknown.

We here systematically re-examine Morgan’s “gradient hypothesis”^8^ in the hope of obtaining new insights into planarian tissue polarity specification. While we were unable to replicate the breakdown of regeneration specificity in narrow cross pieces of the modern model species *S. mediterranea*, we were able to replicate the finding in the *Girardia* genus that Morgan likely worked with^25^. Using a comparative approach, we find that *G. sinensis notum* expression occurs symmetrically at both wound sites and that *notum* expression symmetry contributes to the breakdown of regeneration specificity in very narrow *G. sinensis* cross-pieces. To address how broader *G. sinensis* cross-pieces manage to maintain regeneration polarity nevertheless, we developed a pharmacological approach for the selective perturbation of cWnt gradient shape prior to amputation and show that reductions in gradient steepness increase double-head frequency. With the body size-dependence of double-head formation and the size-dependent decrease in cWnt gradient steepness as additional evidence, our results indicate that the cWnt gradient contributes to tissue polarity in *G. sinensis*. Overall, our results suggest that planarian tissue polarity consists of multiple cues acting in parallel and that differential reliance on these cues contributes to the species-dependence of regeneration defects.

## Results

### Comparative analysis of regeneration specificity

Inspired by Morgan’s classical experiments^6,7^, we first set out to replicate the reported breakdown of regeneration specificity and regeneration of double-headed animals in very narrow cross-pieces in the modern model species *S. mediterranea* (Fig. 1A). Initial experiments established a minimal piece length of ∼1 mm that is required for successful regeneration in this species, with narrower pieces failing to seal the wound and disintegrating soon after cutting in our hands (data not shown). We started with large *S. mediterranea* of 18 mm body length to minimise the piece length relative to body length. The 1 mm or 2 mm long cross-pieces that we examined consequently correspond to 1/18th or 1/9th of the original body length (Fig. 1B-C). To our surprise, we failed to observe even a single double-headed regenerate across two experimental series and ∼400 individual pieces. Almost all of the pieces correctly established regeneration specificity irrespective of piece length, A/P axis position origin or body size, with only a very small proportion of pieces (< 5%) displaying failures of head or tail regeneration (Fig. 1B-E). While we cannot exclude the possibility of length-dependencies in *S. mediterranea* regeneration specificity below the piece length limits required for piece survival, our results stress the fascinating robustness of regeneration specificity in *S. mediterranea* that was one of the selection criteria as model species^26^. Nevertheless, the fact that the rare head regeneration failures all occurred on pieces of anterior origins might hint at slight position-dependencies even in *S. mediterranea* regeneration.

**Figure 1:**
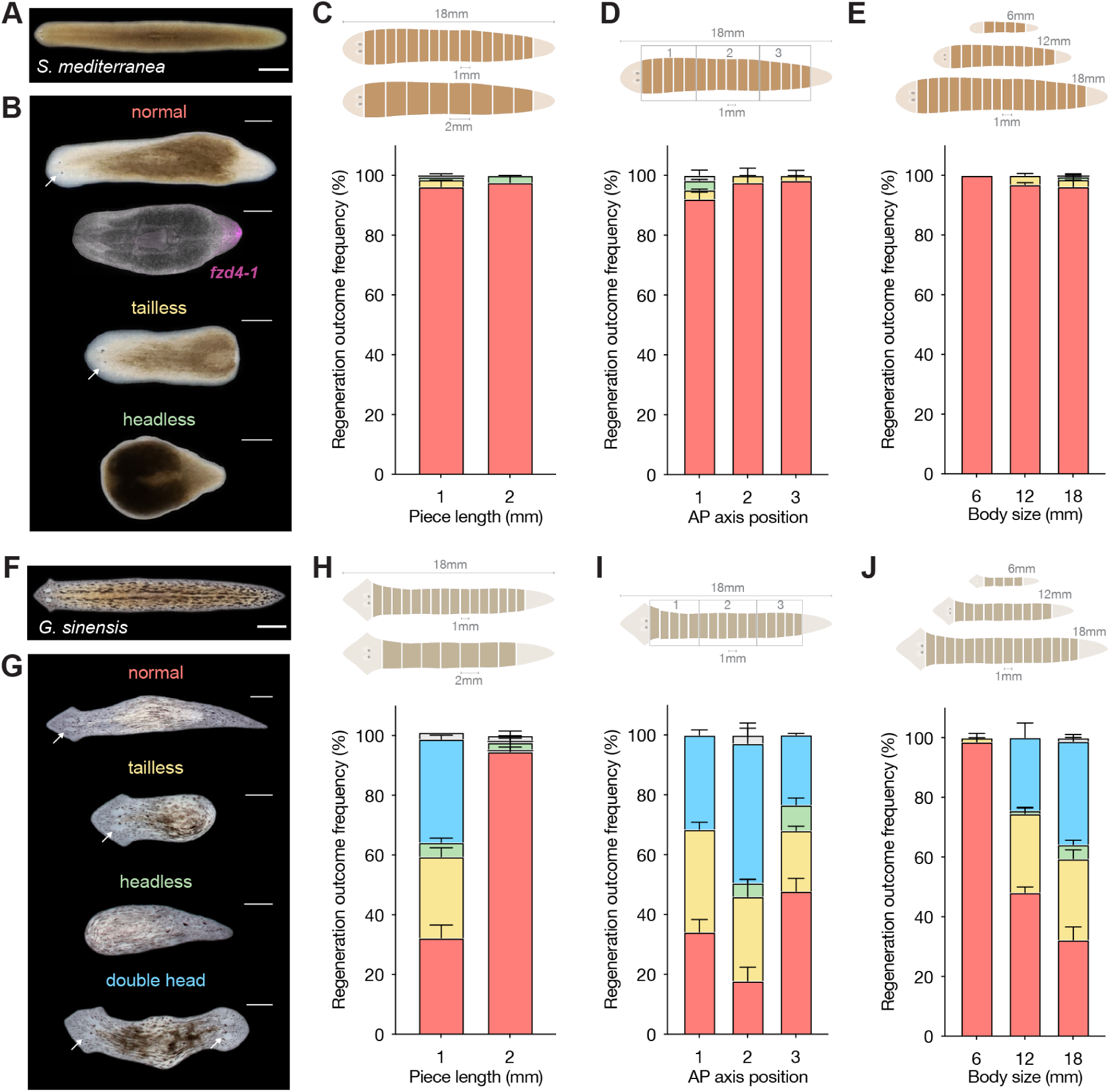
Comparative analysis of head/tail regeneration specificity. A) Uninjured *S. mediterranea*. Scale bar: 2000 µm. B) Representative live images of scored regeneration outcomes in *S.mediterranea* transverse pieces. The expression of the tail marker gene *fzd4-1* by fluorescent whole mount in situ hybridization (FISH) is additionally shown for the “normal” phenotype. White arrows indicate eye spots as morphological head marker. Scale bars: 500 µm. Scoring of regeneration outcomes 14 days post-amputation (dpa), depending on C) piece length, D) A/P axis origin or E) body size. Cartoons illustrate amputation paradigms, colours regeneration outcomes as colour-coded in (B). Grey stacks in plots represent pieces with a regeneration phenotype that could not be unambiguously assigned to one of the four categories. n = 394 pieces. N = two independent experiments. Error bars represent the standard error of the mean (SEM). F-J) As in A-E, but for *G. sinensis* and scored at 21 dpa. n = 540 pieces. N = two independent experiments.

Confronted with the discrepancy between our results and Morgan’s, we next considered the possibility of a species-dependence. *S. mediterranea* is a European species, while T.H. Morgan conducted his experiments at the Marine Biological Laboratory in the USA and worked with a locally sourced species at the time referred to as *Planaria maculata*^6,7^. Reasoning that Morgan’s *P. maculata* likely corresponds to the present-day designation *Girardia tigrina*^25^, we screened several representatives of the *Girardia* genus that we maintain as part of our planarian species live collection^14^ and identified one that we identified as a strain of *Girardia sinensis* (Supp. Fig. 1)^27^ that frequently regenerates double-heads (Fig. 1F). We observed that 36% of 1 mm cross-pieces cut from 18 mm long worms regenerated double-heads, while 2 mm cross-pieces cut from same sized cohorts regenerated predominantly normal animals (Fig. 1G-H). Therefore, these results both confirm Morgan’s findings of length-dependence of regeneration specificity in the genus *Girardia* and, in addition, reveal clear species-dependencies in planarian regeneration specificity. Encouraged by this finding, we next queried the position-dependence of regeneration specificity in *G. sinensis*. Interestingly, we found that 1 mm cross-pieces from the trunk region displayed the highest proportion of double-headed regenerates and other regeneration error categories, with only < 20% of pieces regenerating normal animals (Fig. 1I). Therefore, regeneration specificity is generally position-dependent in *G. sinensis* and least robust in the trunk region of the animal. To assay possible body size-dependencies in the piece length-dependence of regeneration specificity in *G. sinensis*, we complemented the previous regeneration scoring of 1 mm cross-pieces in 18 mm (large) animals with equivalent quantifications of 1 mm cross-pieces cut from 12 mm (medium) or 6 mm (small) animals (Fig. 1J). The regeneration of double-heads in 40%, 25% or 0% of 1 mm fragments originating from large, medium or small animals clearly reveals an additional body size-dependence of regeneration specificity in *G. sinensis*, meaning that the probability for double-head regeneration for a given piece length increases systematically with body size. Overall, our results demonstrate the piece length-, position- and body size-dependence of regeneration specificity in *G. sinensis*, which sharply contrasts with the robustness of regeneration specificity in the model species *S. mediterranea*.

### Wound site-symmetric *notum* expression contributes to the reduced *G. sinensis* regeneration specificity

To investigate possible causes for the dramatic species dependence in regeneration specificity between the two species, we first assembled and annotated a comprehensive *G. sinensis* transcriptome using our established PlanMine pipeline^28^. RNAi-mediated knockdown of the *G. sinensis β-catenin-1* homologue resulted in a high penetrance of double-head regeneration, confirming both the broad conservation of cWnt inhibition as necessary and sufficient prerequisite for planarian head specification^11–14^ and the functionality of the RNAi pathway in *G. sinensis* (Supp. Fig. 2A). In *S. mediterranea*, the asymmetric expression of the Wnt antagonist *notum* is responsible for cWnt inhibition at sites of head regeneration^21^. In regenerating *S. mediterranea* 2 mm trunk pieces cut from 6 mm animals, *notum* expression was induced predominantly at anterior-facing wounds as early as 6 h post amputation (Fig. 2A), as previously reported^21^. Although some cells in the vicinity of posterior wounds also induce *notum* expression, the majority of *notum*-expressing cells were located within 600 µm of anterior wound sites and the maximum asymmetry in terms of expressing cells was reached by 18 h (Fig. 2B-C), consistent with previous results. In 2 mm *G. sinensis* trunk pieces cut from 6 mm animals, the single *notum* homologue was also wound-induced with similar kinetics and in a similar salt-and pepper pattern that reflects muscle cell expression in *S. mediterranea*^29^(Fig. 2D). Strikingly and in sharp contrast to *S. mediterranea*, the number of *notum* expressing cells was nearly identical between anterior and posterior wounds without any discernible A/P asymmetry at any of the examined time points (Fig. 2E-F). High-resolution imaging further confirmed the stark difference between the two species, with barely detectable versus clear *notum* expression at equivalent posterior positions of the two species at peak expression (18 hpa; Fig. 2G-H). Moreover, the invariance of *notum* expression symmetry across all pieces (Fig. 2F) and the fact that the examined 2 mm *G. sinensis* pieces invariably establish correct regeneration specificity (Fig. 1J) imply that the posterior wound expression is not a consequence of already disturbed tissue polarity. Therefore, our results indicate that *G. sinensis* lacks *notum* expression asymmetry, at least in terms of detectable *notum*-expressing cells, and that *notum* expression asymmetry is not necessary for head/tail regeneration specificity in this species.

**Figure 2:**
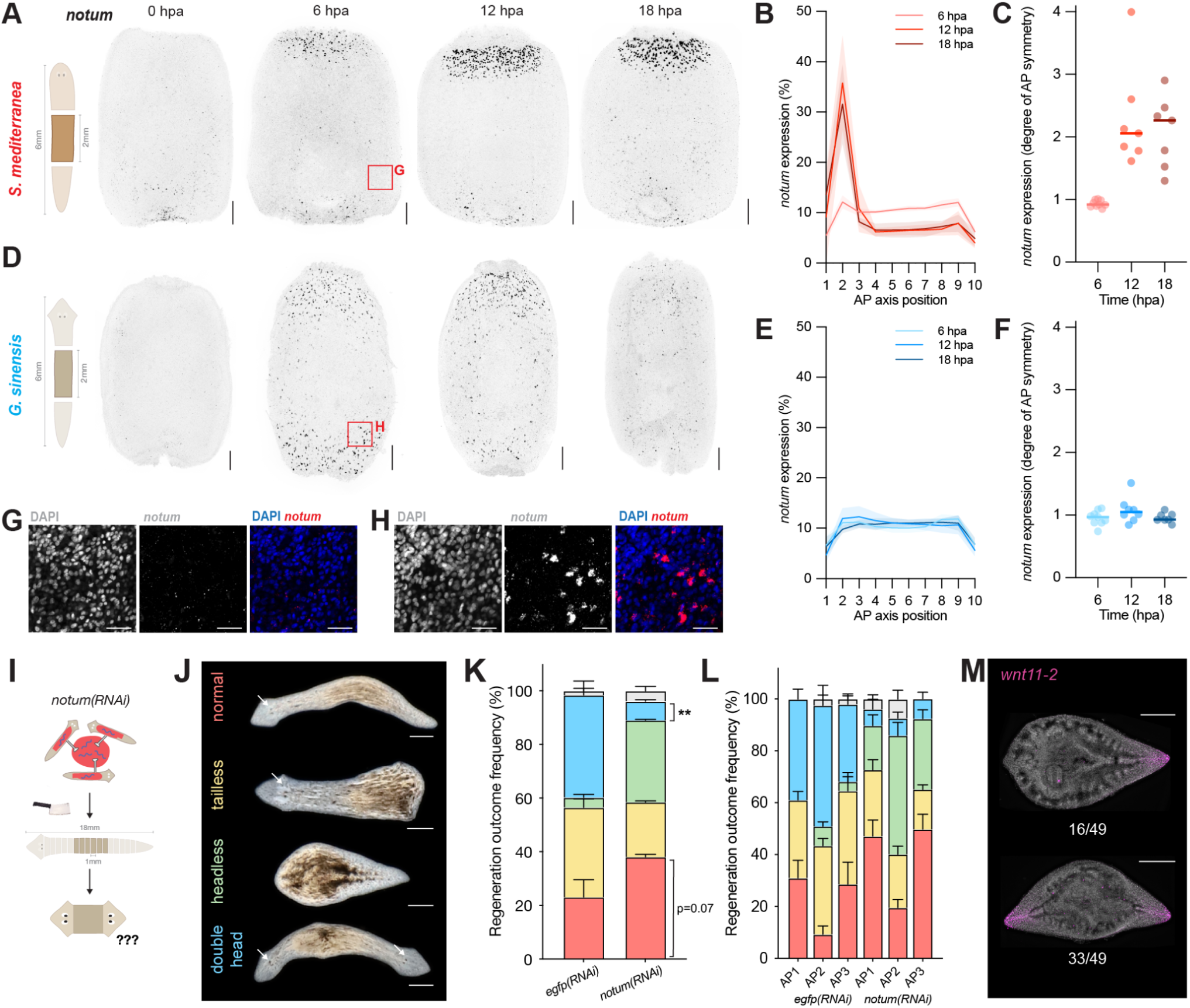
Wound site-symmetric *notum* expression contributes to the reduced *G. sinensis* regeneration specificity. A) Whole-mount *notum* fluorescence in situ hybridisation (FISH) time course on trunk pieces from 6 mm long *S. mediterranea*, as cartooned. Images represent inverted grayscale, maximum intensity projections of Z-stacks from the surface of the specimen to the dorsoventral boundary. hpa: hours post-amputation. Scale bars: 200 µm. B) Quantification of *notum* fluorescence signal distribution along the A/P length of the regenerating pieces (% of total signal located in each equidistant axial bin) at the indicated time points. Error bands represent standard deviation (SD). C) Degree of symmetry (anterior half/posterior half) derived from the data shown in (B). Each point represents one individual piece, the horizontal line the average/median. N = one representative experiment D-F) As in (A-C), but for *G. sinensis*. G) Zoom-in of boxed region in (A), with DAPI-stained nuclei and *notum* FISH signal shown separately and merged. Each image represents a single 3 µm optical section through subepidermal tissue. Scale bars: 50 µm. H) As in (G), but for *G. sinensis*. I) Scheme of the *G. sinensis notum(RNAi)* experiment. J) Representative live images of scored regeneration outcomes. White arrows indicate eye spots as head marker. Scale bars: 500 µm. K) Quantification of regeneration outcomes in control (egfp) versus *notum(RNAi)*. Same colour-coding as in (J). Grey stacks represent pieces with a regeneration phenotype that could not be unambiguously assigned to one of the scored outcomes. Error bars represent SEM. n = 792 pieces. N = three independent experiments. ** p < 0.01, as assessed by Sidak multiple comparisons test. L) The same data as in (K), but additionally segregated according to the A/P axis origin of the pieces (AP1 = anterior third; AP2= central third; AP3= posterior third). M) Whole-mount FISH with the tail marker *wnt11-2* on headless *G. sinensis* pieces. Representative images of pieces with respectively one and two *wnt11-2* expression domains and their relative frequency are shown. n = 49 pieces. N = one representative experiment. Scale bars: 500 µm.

To ask whether *notum* nevertheless maintains a role in the determination of regeneration specificity in *G. sinensis*, we performed RNAi by feeding and examined the consequences of *notum* knockdown on the regeneration outcome of 1 mm pieces cut from 18 mm animals (Fig. 2I). *egfp(RNAi)* controls displayed the expected 40% of double-heads and 23 % of normal regenerates (Fig. 2J-K), consistent with our previous results for this cutting paradigm (Fig. 1H). *notum(RNAi)* increased the proportion of wild type animals to ∼40%, thus partially “rescuing” the species-specific regeneration defect of *G. sinensis* (Fig. 1H). In addition, *notum(RNAi)* significantly reduced the fraction of double-headed regenerates to ∼10%, matched by a corresponding increase in the fraction of “headless” regenerates that appeared to form morphologically normal tails. The effects of *notum(RNAi)* were largely independent of A/P position (Fig. 2L), indicating that the position-dependence of regeneration specificity in *G. sinensis* (Fig. 1I) is independent of *notum*. Examination of the tail marker homologue *wnt11-2* confirmed that the “headless” regenerates indeed form bona-fide tails, including a majority of double-tailed regenerates (Fig. 2M), as in *S. mediterranea notum(RNAi)*^21^. Overall, the increase in tail and decrease in head regeneration are consistent with the conservation of the cWnt inhibitory role of *notum* at *G. sinensis* wound sites and, in conjunction with our expression data (Fig. 2A-H), with a likely permissive *notum* role in *G. sinensis* regeneration specificity. Significantly, the rescue of wild-type regenerates by *notum(RNAi*) suggests that the symmetric *G. sinensis notum* expression contributes to the formation of double-heads and thus to reduced regeneration specificity.

### Conservation of the axial cWnt signalling gradient in *G. sinensis*

The fact that 20% of 1 mm or 100% of 2 mm *G. sinensis* pieces display normal head/tail regeneration specificity despite symmetric *notum* expression implies that polarity cues other than *notum* must mediate regeneration specificity in *G. sinensis*. Given Morgan’s original hypothesis^8^ and the intermittent demonstration of an organismal-scale cWnt signalling gradient along the A/P axis in the model species *S. mediterranea*^16,17^, we decided to explore whether a cWnt signalling gradient might mediate the residual tissue polarity in *G. sinensis*. An initial search for cWnt signalling components in our *G. sinensis* transcriptome revealed a near-identical set of core pathway components as in *S. mediterranea*^11^, with the putative absence of *wnt11-3* as only exception (Fig. 3A; Supp. Table 1). To probe whether the tail to head cWnt signalling gradient was also conserved in *G. sinensis*, we extended our previously established β-catenin-1 quantification by quantitative Western blotting (WB)^16^ to *G. sinensis.* Note that β-catenin-1 levels can provide a useful proxy for planarian cWnt signaling due to the exclusive signalling function of this specific gene homologue^30^ and the demonstrated Wnt-dependence of its detectability in lysates^16^. As our existing anti-β-catenin-1 monoclonal antibody demonstrated a relatively low affinity for *G. sinensis* β-catenin-1^14^, we generated further monoclonal antibodies against the highly conserved N-terminus of β-catenin-1 (Fig. 3B), i.e. the homologue mediating cWnt signalling^30^. One (clone 3588-D84-1) yielded a single prominent band at the expected size of ∼110 kDa that was sensitive to β-catenin-1 knockdown in both *S. mediterranea* and *G. sinensis* (Supp. Fig. 2B). Performing quantitative WB along the A/P axis with our new antibody, we demonstrate that both species deploy a tail-to-head gradient of β-catenin-1 abundance, with the highest levels measured at the tail tip (Fig. 3C). This result, taken together with observations in *Schmidtea polychroa*^17^ and many species across planarian phylogeny^14^ suggest that the β-catenin-1 gradient (hereafter referred to as “cWnt gradient”) may be generally conserved in planaria. Finally, we asked whether the A/P axis gene expression patterns associated with the cWnt gradient in *S. mediterranea* are conserved in *G. sinensis.* Spatially-resolved transcriptome profiling in 10 equidistant sections along the A/P axis as in a previous *S. mediterranea* dataset^16^ revealed that *G. sinensis* broadly recapitulates the axial gene expression patterns of *S. mediterranea*, with prominent clusters of organism-scale anterior-to-posterior and posterior-to-anterior gene expression gradients apparent in our data (Fig. 3D-E; Supp. Table 3). Significantly, the components of the tail cWnt pathway component module and many cWnt signalling target genes expressed in tail gradients^16,31^ were similarly expressed in tail gradients in *G. sinensis* (Fig. 3F). Interestingly, the expression pattern of the *G. sinensis notum* homologue was shifted to the posterior rather than anterior tip as in *S. mediterranea*^21^, consistent with its divergent transcriptional regulation. Overall, our analysis revealed that both the shape and the patterning function of cWnt signalling are broadly conserved in *G. sinensis*, making a role in polarity specification plausible.

**Figure 3:**
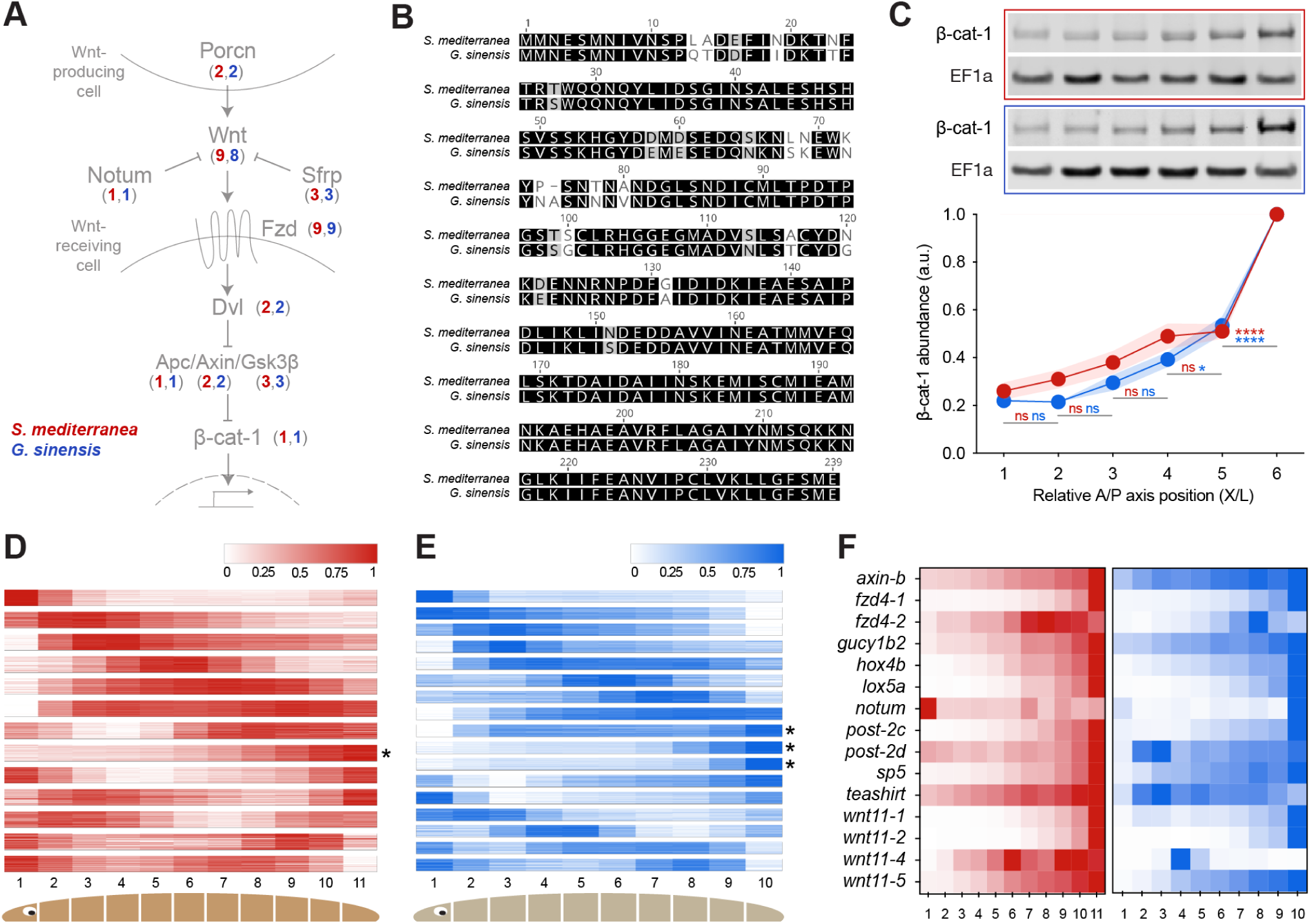
Conservation of the axial cWnt signalling gradient in *G. sinensis*. A) Schematic illustration of the cWnt signalling pathway and number of homologues of the indicated pathway component in *S. mediterranea* (red) or *G. sinensis* (blue). B) Alignment of residues 1-238 of *S. mediterranea* and *G. sinensis* β-catenin-1 that were used as the antigen for antibody production. Black, grey and white highlighting indicate amino acid conservation, conservative replacement and non-conservative replacement. B) Quantitative Western blot analysis of β-catenin-1 abundance as proxy for cWnt signalling activity along the A/P-axis in *S. mediterranea* and *G. sinensis*. 6 mm long specimens were cut into 6 equiproportional transverse pieces that were individually blotted; 1 = head piece; 6 = tail piece. Traces represent the average of four independent experiments (each with >12 individuals per species) and error bands the SEM. β-catenin-1 quantifications were normalised to the loading control in each lane (EF1a) and 6-point measurement sets were further normalised to their p6 sample. ns = not significant, p > 0.05; * p < 0.05, **** p < 0.0001, as assessed by Tukey multiple comparisons testing on adjacent positions (e.g. p5 vs p6). D-E) Clustering of gene expression profiles along the A/P axis of *S. mediterranea* (D) and *G. sinensis* (E), determined by RNAseq of specimens cut into 10 (*G. sinensis*) or 11 (*S. mediterranea*) equiproportional tissue pieces after pharynx removal. Colour scales represent the maximum-normalised expression of each gene. Asterisks indicate posterior-to-anterior gradient clusters. Source data are provided in Supp. Table 3. Plotted values represent a pool of 10 individuals (D) or the average of four individuals (E). The data used to generate (D) were previously reported^16,48^. F) Expression of putative cWnt target genes along the A/P axis of *S. mediterranea* (red) and *G. sinensis* (blue). *S. mediterranea* genes were selected from two published studies^16,48^.

### Establishment of a pharmacological approach for time-resolved cWnt gradient manipulations

If the cWnt gradient were indeed a polarity cue, then the experimental flattening of the gradient prior to amputation should reduce the difference in signalling activity between the two wounds and thus affect head/tail regeneration specificity. However, RNAi as the standard loss of function approach in the field is unsuitable for this purpose, as many of the signalling pathway components that shape the cWnt signalling gradient in intact animals also mediate critical post-amputation cWnt signalling dynamics during head/tail specification in regenerating pieces^16,32^. Pharmacological cWnt signalling pathway interventions might offer the requisite temporal control to separate pre- and post-amputation cWnt signalling; however, pharmacological modulation of cWnt signalling has not been demonstrated in planarians to date. We therefore screened a total of 20 commercially-available small molecule cWnt signalling inhibitors and activators in the model species *S. mediterranea* (Supp. Fig. 2C), with regeneration as functional readout. Only treatments with one of the compounds, Wnt-C59 (hereafter “C59”), produced a dose-dependent appearance of double-head regenerates characteristic of cWnt loss-of-function^18–20^, reaching up to 40% at 50 µM (higher concentrations were not possible due to C59 solubility limitations; Fig. 4A-B). C59 inhibits cWnt signalling by targeting the membrane-bound O-acyltransferase Porcupine (Porcn) required for post-translational modification, secretion and activity of Wnts^33^. Consistent with Porcn as the target of C59 in planaria, we found that RNAi of a so far uncharacterized *porcupine* homologue (designated here as “*porcn-b*”) caused double-head regeneration – and thereby phenocopied C59 – in around 25% of *S. mediterranea* experimental specimens (Fig. 4C-D). The broad expression of *porcn-b* across cell types in single-cell RNAseq datasets^34^ contrasts with the known gut-specific expression of *porcn-a* in *S. mediterranea*^31,35^, and is compatible with a functional role of *porcn-b* in the secretion of Wnt ligands from muscle cells (Supp. Fig. 2D). Encouraged by the results in the model species, we next examined the efficacy of C59 treatments in *G. sinensis* (Fig. 4E-F). Interestingly, C59 treatments caused 100% double-head regenerates already at a five-fold lower dose (10 µM), indicating that *G. sinensis* is more sensitive to C59 than *S. mediterranea* (Fig. 4E-F).

**Figure 4:**
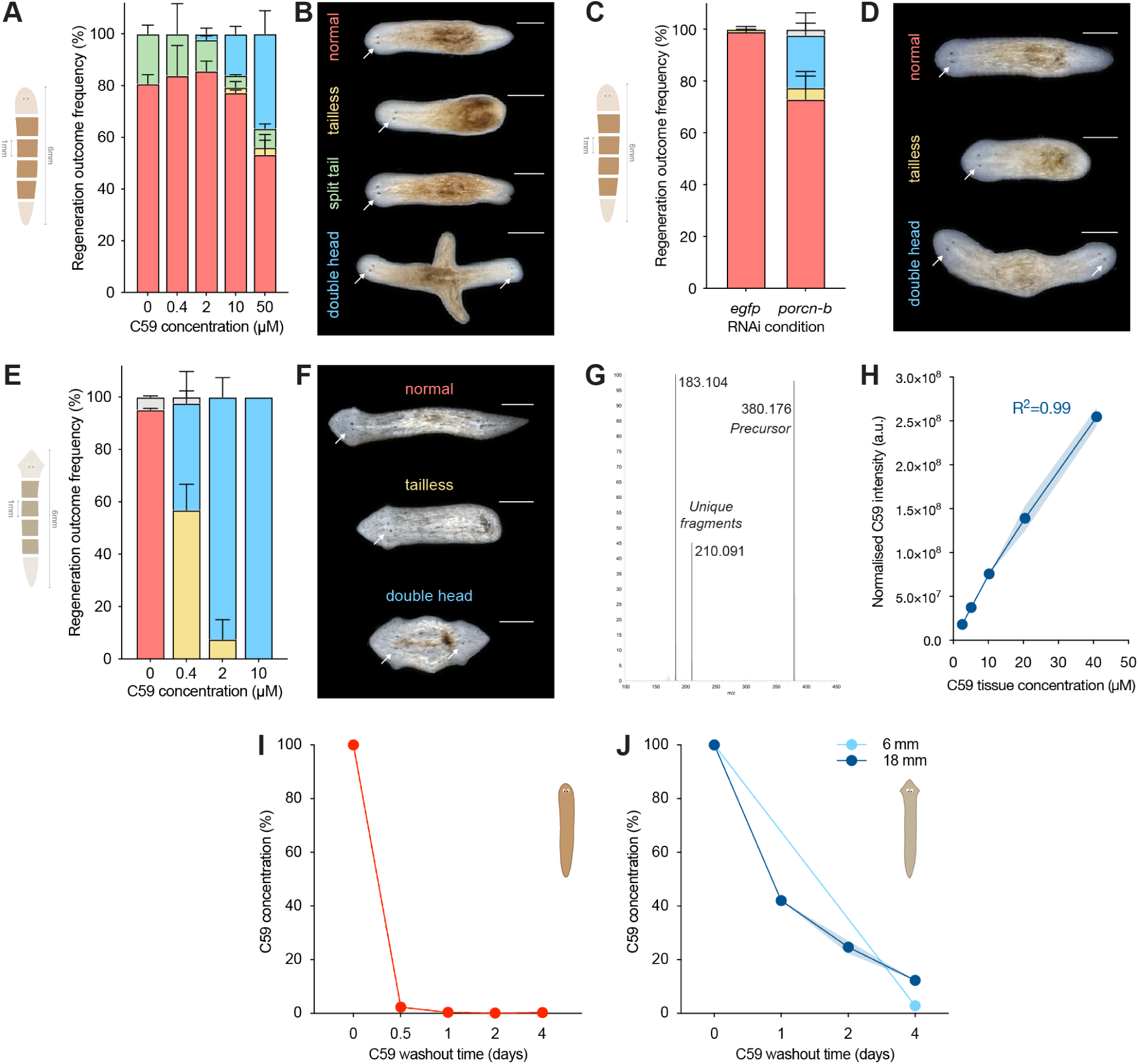
Establishment of a pharmacological approach for time-limited cWnt gradient manipulations. A) Quantification of C59 treatment effects on *S. mediterranea* regeneration outcomes 14 days post-amputation (dpa). The indicated transverse pieces (cartoon) were treated immediately after amputation with the indicated C59 concentrations for 48 h and then left to regenerate in the absence of drug. Stack colors correspond to B), Representative images of the scored regeneration outcomes plotted in (A). White arrows indicate eye spots as head marker. n = 206 pieces. N = two independent experiments. C) *porcn-b(RNAi)* in *S. mediterranea* phenocopies C59 treatment. Quantification of regeneration outcomes of the indicated transverse pieces (cartoon) at 14 dpa versus control (*egfp(RNAi))*. n = 181 pieces; N = two independent experiments. Stack colours correspond to the regeneration outcomes shown in (D), representative images of the scored outcomes plotted in (C). E-F) As in A-B, but in *G. sinensis*. n = 156 pieces. N = three independent experiments. G) Fragmentation pattern of C59 after ms-fragmentation on an Orbitrap mass spectrometer: The singly-charged C59 precursor with m/z 380.176 Da generates two unique fragments with a mass of 183.104 Da and 210.091 Da, respectively. H) C59 quantification by mass spectrometry with sub-micromolar sensitivity. Titration curve, obtained by adding C59 standard to untreated *G. sinensis* tissue extracts to the indicated final concentrations. C59 intensity was normalised to the intensity of a second compound, LGK-974, spiked into all samples at the same concentration as an internal control. Error bands represent the SD. I) C59 clearance kinetics from 12 mm *S. mediterranea.* Specimens were treated for 14 days with 50 µM C59 prior to transfer into drug-free culture medium and sampling at the indicated days post wash-out. Residual C59 concentrations were normalized to the t0 tissue extract drug concentration. n = 3 individuals/time point. N = one representative experiment. J) C59 clearance kinetics from 6 mm and 18 mm *G. sinensis.* Specimens were treated for 7 days with 2 µM C59 prior to transfer into drug-free culture medium and sampling at the indicated days post wash-out. n = 3 individuals per size/time point combination. N = two (6 mm) or three (18 mm) independent experiments. Error bands represent the SEM. Scale bars: 500 µm.

For the intended selective cWnt gradient shape alterations without compromising post-amputation cWnt signalling dynamics, the assessment of the C59 clearance kinetics from planarian tissues further became necessary. We found that C59 can be robustly detected by mass spectrometry (Fig. 4G) and that C59 quantifications in our optimised planarian extract conditions afforded linear detection, even at sub-micromolar concentrations (Fig. 4H). We therefore used mass spectrometry to quantify the residual C59 amounts in extracts of intact *S. mediterranea* and *G. sinensis* at different time points post-treatment. In 12 mm *S. mediterranea*, C59 was rapidly cleared and essentially undetectable already after 12 hours post washout (Fig. 4I). The *G. sinensis* clearance kinetics were significantly slower, respectively achieving 88% and 97% clearance in large (18 mm) and small (6 mm) animals after 4 d of washout (Fig. 4J). Drug clearance rates were largely uniform along the A/P axis (Supp. Fig. 2E) and cWnt signalling remained strongly affected even after 4 d of washout in *G. sinensis* (Supp. Fig. 2F). Overall, these results establish C59 as a viable tool for time-limited cWnt signalling perturbations in *G. sinensis*, but with reduced efficacy in *S. mediterranea*.

### The cWnt gradient contributes to regeneration specificity in *G. sinensis*

To test whether the cWnt signalling gradient indeed contributes to regeneration specificity, we first designed species-specific cWnt gradient perturbations that minimise potential effects on post-amputation cWnt signalling. In *S. mediterranea*, treatments with 50 µM C59 for 14 days were necessary to achieve measurable alterations of the cWnt signalling gradient by quantitative Western Blotting (Fig. 5A-B). Even so, measured cWnt signalling levels were only significantly reduced in the tail tip (piece 6), but not in more anterior body regions. Treated animals still appeared overtly normal after 14 days of drug treatment and morphological abnormalities only became apparent upon even longer treatment intervals (data not shown). Since the rapid C59 clearance kinetics in *S. mediterranea* (Fig. 4I) likely limit drug effects to the first hours post-amputation, we omitted a dedicated wash-out step from the *S. mediterranea* C59 treatment protocol (Fig. 5A). Unlike the C59 treatments during the first 48 h of regeneration (Fig. 4A-B), the C59 pre-treatment of 6 mm *S. mediterranea* did not affect the regeneration specificity of 1 mm cross-pieces (Fig. 5C-E). Therefore, at least the modest gradient shape alterations achievable by our treatment protocol did not affect regeneration specificity in *S. mediterranea,* and the double-head formation in Fig. 4A were likely the result of post-amputation cWnt signalling inhibition. In the case of *G. sinensis*, C59 pre-treatments with 2 µM C59 for 7 days were already sufficient for achieving a significant reduction of cWnt signalling levels at the tail tip, again indicating a greater treatment efficacy in this species (Fig. 5F-G). In order to account for the slower clearance kinetics in *G. sinensis*, we incorporated a 4 day wash-out interval prior to amputation (Fig. 5F), which is sufficient for near-complete C59 clearance in 6 mm animals (Fig. 4J). Although some recovery of the cWnt signalling gradient occurred during the wash-out interval, the tail tip signalling levels and hence the slope of the cWnt signalling gradient in the tail remained significantly reduced at the time of amputation (Supp. Fig. 2F). Intriguingly and in contrast to *S. mediterranea*, the C59 pre-treatment of 1 mm cross pieces from 6 mm animals induced ∼10% of double-headed regenerates and further increased the fraction of “tailless” animals, while DMSO-treated controls displayed the high regeneration specificity previously demonstrated for 1 mm cross-pieces from small animals (Fig. 5H vs. 1J). Moreover, the regeneration specificity effects were location-dependent, with double-heads originating exclusively from the central body region or the tail of pre-treated donor animals (Fig. 5I-J). Given the near-complete clearance of C59 at the time of amputation in 6 mm animals (Fig. 4J), these results were therefore consistent with an instructive role of pre-amputation cWnt signalling in regeneration specificity and thus a role of the cWnt signalling gradient in determining regeneration specificity in *G. sinensis*.

**Figure 5:**
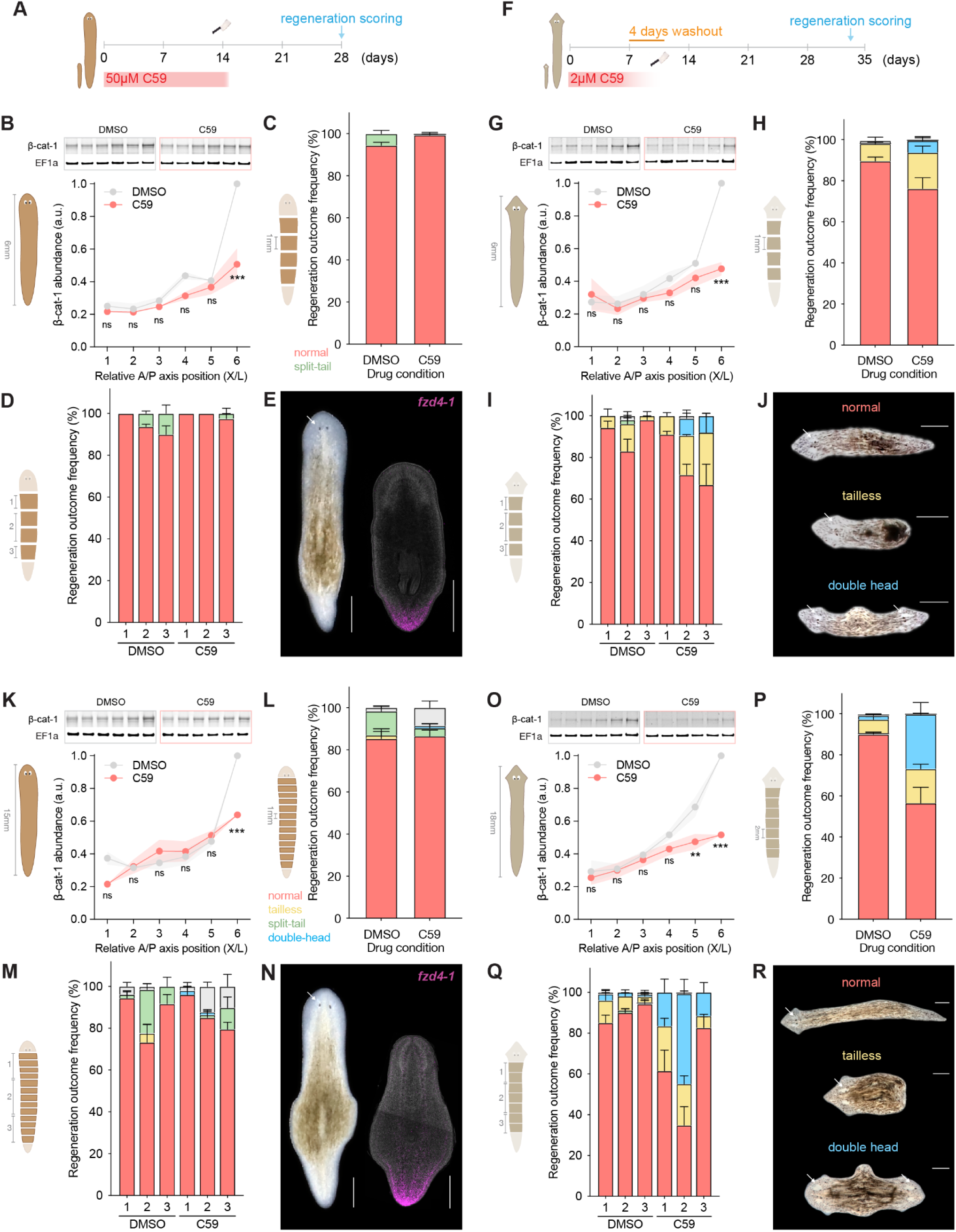
The cWnt gradient contributes to regeneration specificity in *G. sinensis*. A) Cartoon of the c59 treatment protocol used for testing cWnt gradient contributions to *S. mediterranea* regeneration specificity. B) Western blot analysis of β-catenin-1 abundance/cWnt signalling activity along the A/P axis in the indicated 6 mm long *S. mediterranea*, after 14 days pre-treatment with 50 µM C59 or vehicle (0.75% DMSO). The β-catenin-1 signals in lysates of equiproportional transverse pieces of the indicated specimens were normalized to the loading control (Smed-EF1a) and each 6-point measurement set was further normalised to the tail piece (p6) of the DMSO-control. n = 10 individuals per group. N = three independent experiments. Error bars represent the SEM. C) Regeneration outcome quantification of the indicated *S. mediterranea* transverse pieces under the same vehicle/C59 treatment protocol as in (A, B). Colour-coding of regeneration outcomes as indicated. n = 315 pieces. N = four independent experiments. D) As in (C), but with pieces separated into three A/P axis bins as cartooned. E) Representative live image (left) and tail marker *fzd4-1* FISH image (right) of “normal” regenerated fragments, confirming wild-type regeneration in this experiment. F-I) As in (A-D), except with *G. sinensis*. Note the lower drug concentrations (0.5% DMSO vehicle or 2 µM C59), shorter pre-treatment and 4 days of subsequent drug washout for these experiments. N = 195 pieces across three independent experiments (H-I). J) Representative live images of regeneration outcomes, colour-coded as in H-I K-N) As in (B-E), but with 15 mm long *S. mediterranea*. n = 3 individuals/group (K) and 373 pieces (L-M). N = 3 independent experiments (K-M). O-R) As in (B-E) but with 18 mm long *G. sinensis*. n = 3 individuals/group (O) and 699 pieces (P-Q). N = five independent experiments (O-Q). ns not significant, p > 0.05; ** p < 0.01; **** p < 0.0001; as assessed by 2-way ANOVA with Bonferroni multiple comparisons test. White arrows indicate eye spots as head marker. Scale bars: 500 µm.

We next examined the effect of C59 pre-treatment in large animals (15 mm long *S. mediterranea* and 18 mm long *G. sinensis*), reasoning that the body size-dependence of regeneration specificity (Fig. 1J) might render large animals more sensitive to the mild gradient perturbations that our treatment protocol affords. In *S. mediterranea*, the two-week C59 pre-treatment elicited similarly modest effects on the cWnt gradient shape as in 6 mm animals (Fig. 5K vs. 5B) and 1 mm transverse fragments displayed similarly high regeneration specificity as the DMSO-treated controls (Fig. 5L,N). However, 2 out of 187 fragments examined did regenerate double-heads (Fig. 5L, Supp. Fig. 3A), with both originating from the anterior or central body regions (Fig. 5M). Even though we never observed double-heads amongst an even greater number of untreated controls (Fig. 1C), the low frequency of the phenotype again emphasised the robustness of regeneration specificity in *S. mediterranea*. In the case of *G. sinensis*, the 7d C59 pre-treatment of 18 mm animals resulted in a more significant gradient flattening with our 6-point assay than in 6 mm animals, with significant reductions of cWnt signalling in both pieces 5 and 6 (Fig. 5O). In addition, C59 pre-treatment elicited >15% of double-headed regenerates in 2 mm cross-pieces, which otherwise largely regenerated normal animals in untreated and DMSO treated controls (Fig. 5P,R vs. 1H; Supp. Video 1). The increase in double-headed regenerates is unlikely due to the somewhat slower clearance of C59 from 18- as compared to 6 mm animals (Fig. 4J), as the application of the residual drug concentration to the culture medium during the regeneration interval resulted in fewer double-headed animals than pre-amputation treatments (Supp. Fig. 3B-E). Interestingly, the fragments originating from central body regions were most sensitive to C59 pre-treatments, with >50% of fragments regenerating double-heads (Fig. 5Q). Notably, the regeneration outcome of the C59 pre-treatments 2 mm pieces from 18 mm *G. sinensis* almost perfectly phenocopies that of “wild-type” 1 mm pieces from 18 mm *G. sinensis* in terms of both frequency and position-dependence of double-heads (Fig. 5P-Q vs. 1H-I). Overall, these results are consistent with a contribution of the cWnt gradient to regeneration specificity in *G. sinensis*, that is a threshold-level difference in cWnt signalling activity across a piece as requirement for differential head/tail regeneration.

### cWnt gradient scaling

If *G. sinensis* tissue pieces were indeed using the steepness of the cWnt gradient (referred to as “slope” in the following) as a polarity cue, a progressive flattening of the gradient with increasing body size might explain the body size-dependence of both the critical piece length and C59 efficacy (Fig. 6A). To test this prediction, we examined the scaling of the cWnt gradient in *G. sinensis* and *S. mediterranea*. Using our qWB assay, we quantified β-catenin-1 amounts in 6 equiproportional transverse pieces (i.e., each piece accounting for ⅙ of the length of the A/P axis) of 6 mm, 12 mm and 18 mm long individuals. Plotting the individual measurements by their relative A/P axis position-of-origin revealed remarkably similar shapes of the length-normalised cWnt signalling profile shape between the two species and across the examined body size range, thus clearly indicating that the planarian cWnt signalling gradient scales (Fig. 6B-C). However, in considering the absolute A/P axis length variations (6 mm, 12 mm and 18 mm) and the slight decrease in absolute gradient peak amplitude (tail piece) towards larger body sizes, our data indicate that the slope of the cWnt signalling gradient becomes progressively less steep with increasing body length, thus reducing the difference in signalling activity per A/P axis unit length. The scaling of the cWnt signalling gradient is therefore consistent with the observed size-dependent decrease in regeneration specificity and the greater sensitivity to mild slope reductions in large animals. Overall, the size-dependency of cWnt signalling therefore provides further experimental evidence for a functional contribution of the cWnt gradient slope to tissue polarity specification in *G. sinensis*.

**Figure 6:**
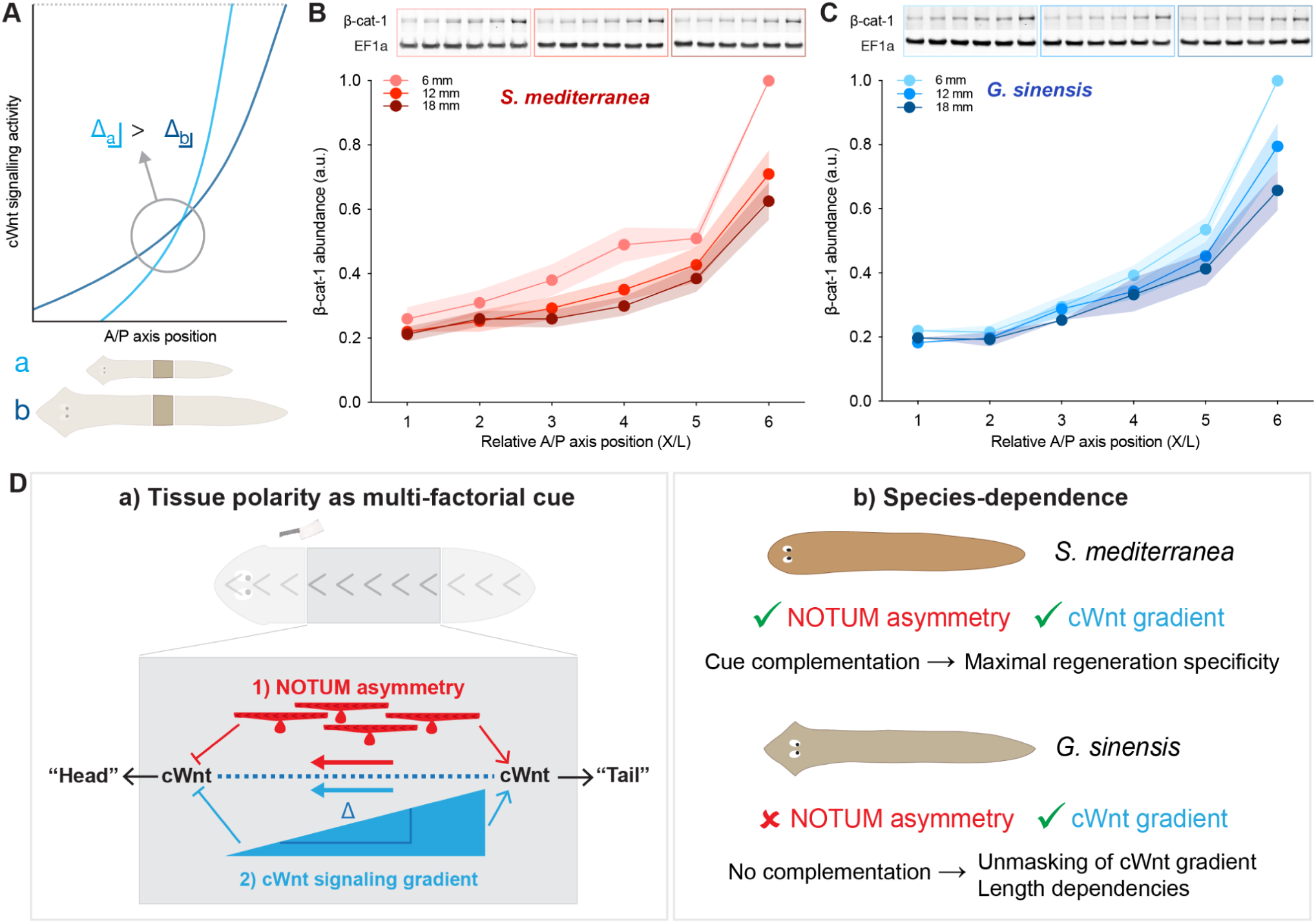
Scaling of the cWnt gradient and implications for planarian tissue polarity. A) Schematic illustration of how cWnt gradient scaling and the associated slope decrease might account for the body size-dependence of regeneration specificity in *G. sinensis* B-C) Western blot quantification of axial β-catenin-1 abundance in 6 mm, 12 mm and 18 mm long *S. mediterranea* (B) and *G. sinensis* (C). The β-catenin-1 signals in lysates of equiproportional transverse pieces were normalised to the loading control (Smed-EF1a) and each 6-point measurement set was further normalised to the tail piece (p6) of the DMSO-control. Traces represent the average of four independent experiments (each with 4-16 individuals per species) and are shown normalised by body length (L). Error bands represent the SEM. ns = not significant, p > 0.05; * p < 0.05, **** p < 0.0001; as assessed by Tukey multiple comparisons testing on adjacent positions (e.g. p5 vs p6). Note that the 6 mm data shown here are the same data as shown in Fig. 3C. D) Working model of the mechanisms underlying planarian tissue polarity (a) and species-dependent regeneration specificity (b). See text for details.

## Discussion

Overall, our results suggest the following model (Fig. 6D): The intrinsic tissue polarity that instructs planarian head/tail regeneration specificity is a multi-factorial cue comprised of at least two distinct pathways: 1) the slope of the organismal cWnt signalling gradient (and possibly other signalling gradients; see below); and 2) asymmetric/anterior-enriched *notum* expression and its upstream regulatory mechanisms. Both cues are capable of independently influencing the autocatalytic and mutually exclusive up- or down-regulation of cWnt signalling at wound sites at 18 h post amputation that signifies the head/tail regeneration commitment^16^. Planarian species in which both pathways are functional (e.g. *S. mediterranea*) display the most robust regeneration specificity due to the functional complementation of both cues. In *G. sinensis*, the loss of anterior-enriched *notum* expression leaves the cWnt gradient as the remaining polarity cue and concomitantly uncovers its intrinsic functional limitations, e.g., piece length-, body size- and position-dependence. Our model provides a rationale for the species-dependent regeneration defect in *G. sinensis* and why Morgan’s seminal demonstration of double-head regeneration from narrow cross pieces^6,7^ could never be replicated in the modern model species *S. mediterranea.* The cWnt gradient slope as polarity cue and the requirement for a minimal activity threshold difference across regenerating cross pieces are consistent with the general length (piece width) dependence of *G. sinensis* regeneration specificity, with our findings of body-size-dependent reductions in cWnt gradient slope and the body size-dependence of regeneration specificity in *G. sinensis* as further confirmation of the model. Finally, our demonstration that experimental reductions in the cWnt gradient slope prior to amputation increase the length-dependence of regeneration specificity provide direct evidence for the contribution of the cWnt signalling gradient to tissue polarity specification in planarians.

Although the concept of gradient-based polarity specification was already proposed by T.H. Morgan more than 100 years ago^8^ and the existence of long-range signalling gradients deployed by the shell-like planarian body wall musculature^29^ have meanwhile been demonstrated at least for the tail-to-head cWnt signalling gradient^16,17^, the possibility of gradient-mediated polarity specification has not featured prominently during the molecular revival of planarian research. Besides the lack of demonstrable piece-length dependence in *S. mediterranea*, part of the reason is the discovery of the critical symmetry-breaking function of asymmetric *notum* expression in *S. mediterranea*^21^. Although the anterior-enriched expression of *notum* in *S. mediterranea* requires cWnt signalling (i.e. the much-reduced expression of *notum* after *β-catenin-1(RNAi)*), the global activation of cWnt signalling by *APC(RNAi)* does not abrogate the anterior-specificity of *notum* expression^21^. Hence, the anterior-specificity of *notum* expression is thought to be independent of the cWnt gradient and although the underlying mechanisms remain poorly understood, recent results point towards the orientation and intrinsic polarisation of the *notum*-expressing longitudinal body wall muscle fibres via *activin-2* and non-canonical Wnt signalling^23,24^. In this context, our findings are important for the following reasons: First, although we cannot formally exclude *notum* expression asymmetry below the detection threshold of our in situ hybridization approach or post-transcriptional asymmetry generation in *G. sinensis* (e.g. anterior-specific translation), our results suggest that *notum* expression asymmetry is not universally conserved in planarians and that concomitantly head/tail specification is possible without *notum* expression asymmetry in cross-pieces above a critical length (e.g. >1 mm in 6 mm animals or >2 mm in 18 mm *G. sinensis*). Second, our demonstration that pre-wounding manipulation of the cWnt signalling profile influences regeneration specificity and that the body size and position-dependencies of regeneration specificity correlate with the measured shape and scaling of the cWnt signalling gradient provide experimental evidence for a polarity-specifying role of this signalling gradient.

Given that the planarian body plan is constitutively polarised via tail-to-head Wnt ligand and receptor expression gradients^16^, and that tail/head regeneration requires activation or inhibition of cWnt signalling at the wound site, the reinforcement of residual cWnt signalling differences across the regenerating piece provides an attractive potential mechanism for ensuring head/tail regeneration in alignment with wound orientation. The known propensity of certain classes of self-organised signalling patterns to reinforce existing spatial differences in signalling activity *in silico*^36,37^, and the intrinsic pattern length scales arising from the diffusion/degradation kinetics of molecules^38^, provide a conceptual framework for envisaging gradient-mediated polarity specification at a mechanistic level. The many transcriptional and post-transcriptional feedback loops in planarian cWnt signalling, and the evidence for mutual antagonism between cWnt-mediated cWnt signalling and a yet-to-be-identified signal patterning the oppositely oriented head-to-tail gene expression gradients^39,40^ as the molecular basis of the tail/head decision^16^, make this view plausible. Furthermore, the low amplitude of both gradients in the central body regions correlates with the low point of regeneration specificity (Fig. 1I), thus also providing a potential explanation for the position-dependence of *G. sinensis* regeneration specificity. In general, the head/tail specification process during planarian regeneration has interesting analogies to symmetry-breaking mechanisms in early embryos, which often involve the context-dependent triggering of self-amplifying but mutually antagonistic signalling networks^41,42^. Furthermore, the deep phylogenetic conservation of symmetry-breaking signalling patterns (e.g. cWnt, Bmp4/Admp) contrasts with the apparently rapid divergence of the upstream mechanisms that trigger symmetry-breaking^43,44^. The intrinsic tendency of self-organising signalling patterns to amplify small initial differences may contribute to the evolvability of symmetry-breaking mechanisms, since any process leading to a small local change in signalling intensity may be sufficient for symmetry-breaking. Similarly, the asymmetry of *notum* expression and the residual slope of the cWnt signalling gradient can be considered as independent but synergistic modulators of the wound site signalling environment, triggering the mutually exclusive head or tail signalling pattern depending on the polarity of the adjacent tissue. The cWnt signalling gradient as a polarity cue is particularly attractive in this context, as it combines polarity encoding (slope of the spatial signalling profile) and symmetry breaking (temporal transition between alternative signaling states) in one and the same mechanism.

That said, the correlation between the various length dependencies in *G. sinensis* regeneration and the scaling of the cWnt signalling gradient does not prove causality and the causal link between gradient slope reductions and regeneration specificity in Fig. 5 is subject to the limitations of our current pharmacological tool kit. Although our drug clearance rate quantifications (Fig. 4I-J; Supp. Fig. 2E-F and 3B-C) make it unlikely that the increased frequency of double-headed regenerates is caused by residual post-amputation cWnt signalling inhibition, C59 treatments allowed only comparatively minor reductions of the cWnt gradient slope, even in *G. sinensis*. The much reduced C59 efficacy in *S. mediterranea*, even at 25-fold higher dosage (Fig. 5), aligns with various anecdotal indications in the field that the model species may be particularly resistant to drug treatments. Given that C59 targets the porcupine-mediated lipidation and activation of de novo-synthesised Wnt ligands^33^, elucidating the extent by which Wnt ligand deposition/stabilization in the extracellular matrix or potentially partially lipidation-independent Wnt signaling contribute to the slow onset and long lasting effect of C59 treatments constitute interesting questions for future studies. In addition, our model predicts that symmetric *notum* expression in *S. mediterranea* should uncover the residual contributions of the cWnt gradient to regeneration specificity. Knockdown of *activin-2* and both *wnt11-1* and *wnt11-2* together has meanwhile been shown to reduce the asymmetry of *notum* expression and to cause a fraction of double-headed regenerates in *S. mediterranea*^23,24^. Whether the penetrance of these phenotypes display any piece length- or body size dependences has not been examined and provides an interesting avenue for future explorations. Alternatively, it is also possible that *S. mediterranea* has reduced or lost the contribution of the cWnt signalling gradient to polarity specification and instead relies entirely on “hard-wired” muscle fibre polarisation and downstream *notum* asymmetry for polarity specification. Finally, it is important to stress that the gradient hypothesis provides a compelling explanation for piece length- body size- and even position-dependence of regeneration specificity, but that other mechanisms may also give rise to the observed length dependencies. One interesting possibility is the anatomical fine structure of muscle fibres and the body size- or species-dependent scaling of associated length scales, along with the elucidation of the mechanistic basis of asymmetric (*S. mediterranea*) or symmetric (*G. sinensis*) *notum* gene expression. Another possibility is that double-head formation might not reflect ambiguities in the upstream polarity cue as assumed by our model, but rather downstream consequences of initially correctly specified wound site signalling patterns (e.g., a larger and size-dependent Wnt inhibition zone associated with early *G. sinensis* head regeneration). Clearly, the visualisation of spatial and temporal signalling dynamics during regeneration and the development of the requisite tools will be ultimately required for a mechanistic understanding of planarian tissue polarity.

Overall, our results once again highlight the extent of mechanistic divergence between planarian species. Together with the previously reported lack of wound-induced *wnt1* expression in *D. lacteum*^11^, the absence of *notum* expression asymmetry in *G. sinensis* demonstrates a lack of conservation of already two presumed core components of planarian regeneration. On the one hand, this highlights the need for extending mechanistic studies beyond the model species to distinguish species-specific adaptations from core mechanisms. On the other hand, the experimentally demonstrated importance of asymmetric *notum* expression or wound-induced *wnt1* expression in *S. mediterranea* regeneration raises the challenge of understanding how else regeneration remains possible in their absence. In general, redundancy between parallel-acting pathways, as in our model, is an attractive concept for both the robustness of physiological processes and the evolvability of the underlying mechanisms, and planarians as a model taxon provide ample opportunities for studying it.

## Materials and Methods

### Key resources table

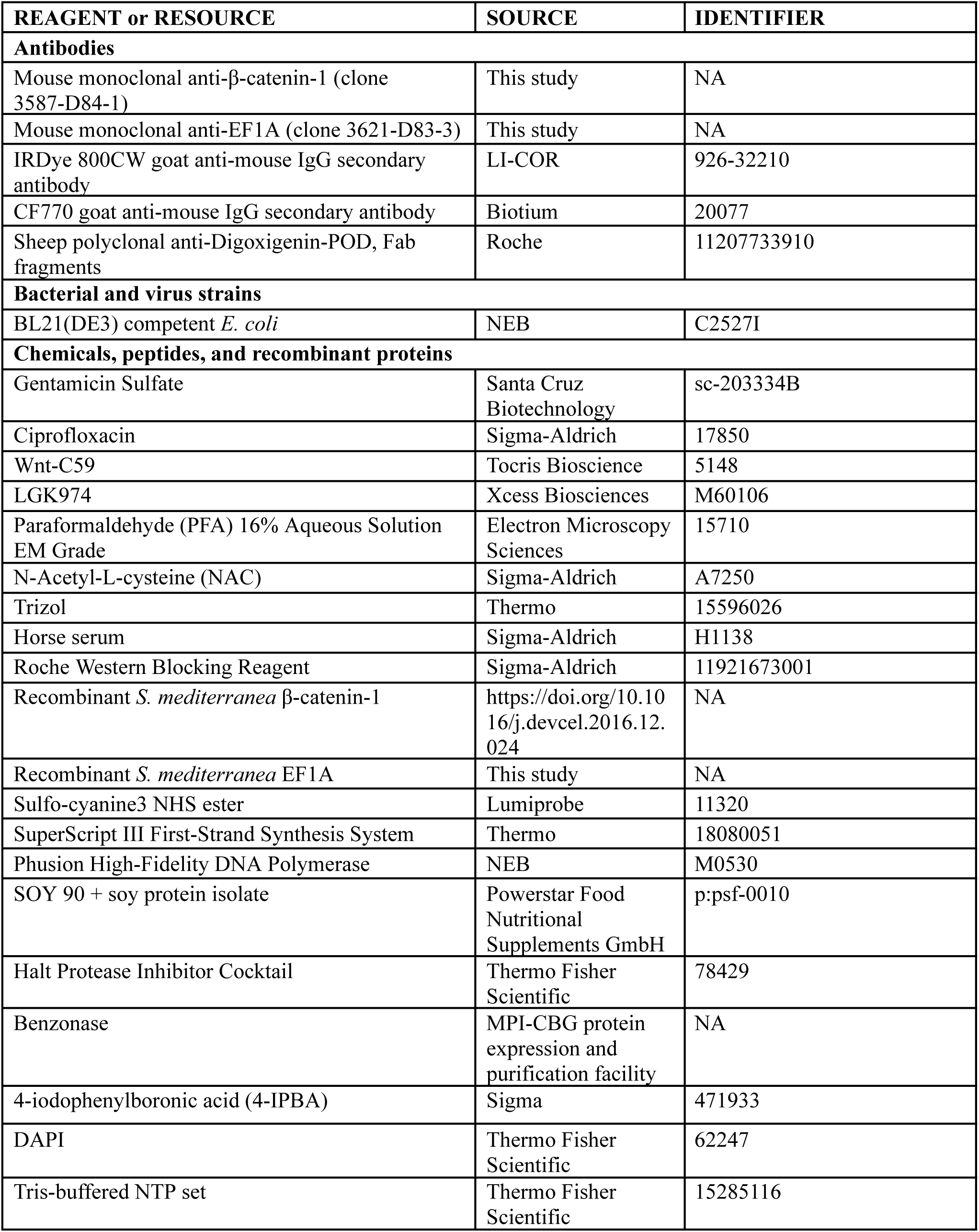
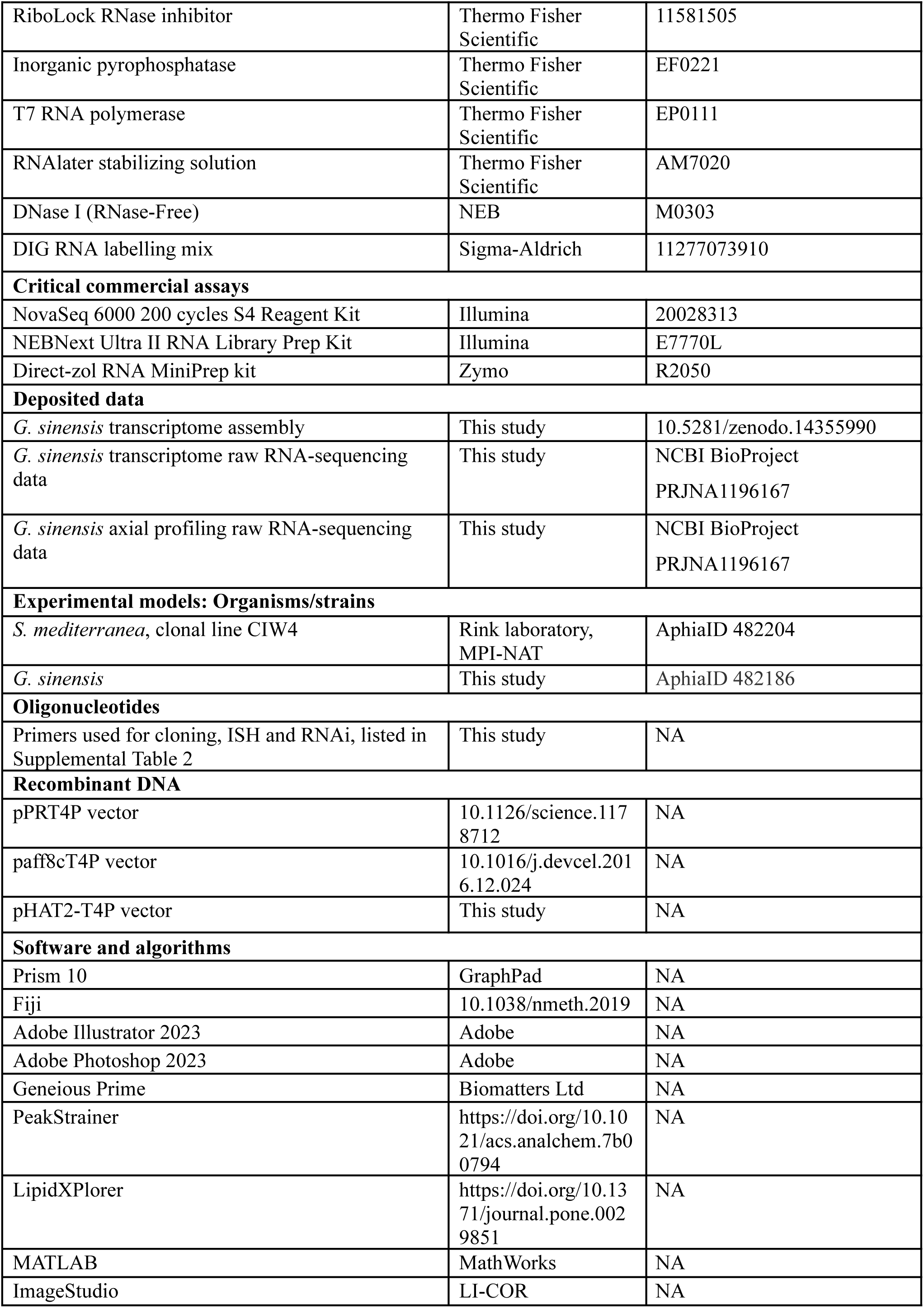
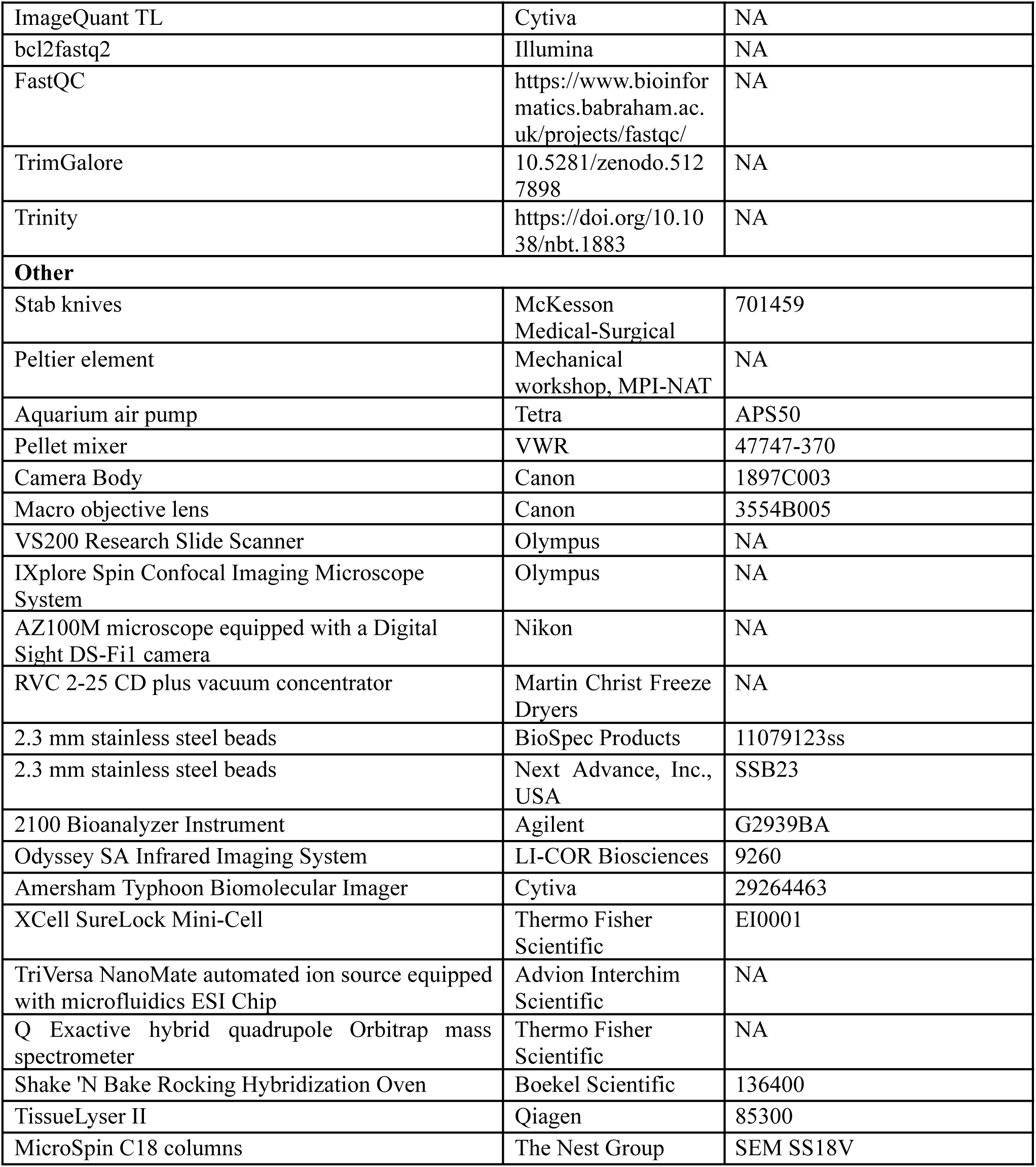

### Resource availability

#### Lead contact

Further information and requests for resources and reagents should be directed to and will be fulfilled by the lead contact, Jochen C. Rink.

#### Materials availability

Materials generated in the course of this study will be made available upon request to the lead contact, Jochen C. Rink.

#### Data and code availability

The previously published *S. mediterranea* transcriptome^11^ is available via PlanMine (https://planmine.mpinat.mpg.de/) and axial RNA-seq data^16^ via NCBI (BioProject PRJNA357536). All raw RNA sequencing reads for the newly reported *G. sinensis* transcriptome and A/P transcriptomics are available via NCBI (BioProject PRJNA1196167). The *G. sinensis* transcriptome assembly is available via Zenodo (10.5281/zenodo.14355990) and will be incorporated into the PlanMine online resource prior to publication. All raw mass spectrometry data will be made available via an online repository prior to publication. All newly reported gene sequences will be submitted to GenBank prior to publication and the accession numbers provided in Supp. Table 2. The code required to reproduce the A/P transcriptomics analysis (Fig. 3 and Supp. Fig. 4) is available via Zenodo (10.5281/zenodo.14356209).

### Experimental models and subject details

*S. mediterranea* (CIW4 strain) and *G. sinensis* (collected near Montpellier, FR) were used for all experiments in this study. *G. sinensis* were maintained in the standard planarian water formulation (1.6 mM NaCl, 1.0 mM CaCl_2_, 1.0 mM MgSO_4_, 0.1 mM MgCl_2_,0.1 mM KCl, 1.2 mM NaHCO_3_)^45^ supplemented with 25 mg/L gentamicin sulfate at 20°C. *S. mediterranea* were generally maintained the same way, except the animals used for the experiments presented in Figure 5, which were kept in water supplemented with 5 mg/L ciprofloxacin instead of gentamicin. Animals were regularly fed organic calf liver paste and starved for at least one week before experiments. Cohorts of the desired body size (6 ± 1, 12 ± 1 or 18 ± 1 mm in length) were isolated by letting animals glide across a petri dish sitting on top of graph paper until they were fully extended and thus measurable.

*S. mediterranea* (CIW4 strain) and *G. sinensis* (collected near Montpellier, FR and maintained in the lab for over 3 years prior to the experiments presented here) were used for all experiments in this study. *G. sinensis* were kept in standard planarian water (1.6 mM NaCl, 1.0 mM CaCl2, 1.0 mM MgSO4, 0.1 mM MgCl2, 0.1 mM KCl, 1.2 mM NaHCO3) containing 25 mg/L gentamicin sulfate, and maintained at 20°C. *S. mediterranea* were generally maintained under the same conditions, but for the experiments shown in Figure 5, they were kept in water supplemented with 5 mg/L ciprofloxacin instead of gentamicin. The animals were regularly fed organic calf liver paste and fasted for at least one week prior to experimentation. To obtain cohorts of the desired body size (6 ± 1, 12 ± 1, or 18 ± 1 mm in length), animals were allowed to glide across a petri dish placed on graph paper until fully extended and measurable.

### Method details

#### Quantitative analysis of regeneration specificity

Animals were immobilised on a chilled peltier element and amputated using a stab knife. Specific-length pieces (e.g., 1 or 2 mm) were isolated through progressive fractionation^46^. For instance, to obtain 1 mm pieces, 18 mm long animals were first cut into three 6 mm segments. Each 6 mm segment was then divided into two 3 mm segments, and each 3 mm segment was further cut into three 1 mm pieces. The first (head tip) and last (tail tip) pieces were excluded due to having only one wound site. The remaining pieces were transferred to well plates or petri dishes with fresh water, which was regularly changed to ensure optimal viability of the regenerating fragments. Regeneration was visually assessed after 14 days for *S. mediterranea* and 21 days for *G. sinensis* using a dissecting microscope. Representative images of various phenotypes are shown in the figures and additional experiment-specific details provided in the respective figure legends.

#### Transcriptome assembly

Total RNA was extracted from pools of both uninjured and regenerating (1, 2, 3, 4, 5, 6, and 7 days post-amputation) sexually mature *G. sinensis* using mechanical homogenization in Trizol, followed by purification with the Direct-zol RNA MiniPrep kit. RNA quality was assessed with a Bioanalyzer, and mRNA was enriched through poly-DT pull-down. Libraries were prepared using standard Illumina mRNA library preparation methods and sequenced on a NovaSeq 6000 with a 200-cycle S4 Reagent Kit. Transcriptome assembly was performed as described previously^47^.

#### Phylogenetic analysis

69 *cox1* sequences with species-level classification^27^ were downloaded from NCBI and imported into the Geneious prime software for phylogenetic comparison to a *cox1* sequence retrieved from the the above-described transcriptome assembly. The phylogenetic tree presented in Supp. Fig. 1 was built using the “Geneious Tree Builder” function with default parameters. Specifically, a distance matrix was built using the “global alignment with free end gaps” alignment type and “65% similarity (5.0/-4.0)” cost matrix, then the tree itself was built using the “Tamura-Nei” genetic distance model, “neighbor-joining” tree build method and no outgroup.

#### Spatially-resolved transcriptomic profiling along the A/P axis

*S. mediterranea* RNA-seq data were generated previously^16,48^. Four 18 mm long *G. sinensis* were dissected into 10 pieces each and immediately placed in pre-chilled RNAlater. Note that *G. sinensis* pharynges were not intentionally removed by sodium azide treatment prior to amputation, but were spontaneously ejected during amputation. Once all pieces were collected, the RNAlater was replaced with 1 ml of Trizol, and the samples were stored at −80°C. After mechanical homogenization with a TissueLyser II (2 minutes at 30 Hz), total RNA was extracted from each of the ten samples using phenol/chloroform extraction, purified with a Direct-zol RNA MiniPrep kit, and assessed for quality with a Bioanalyzer. mRNA was then enriched through poly-DT pull-down. Libraries were prepared using a NEBNext Ultra II RNA Library Prep Kit and sequenced as 100 bp paired-end reads on a NovaSeq 6000 with a 200-cycle S4 Reagent Kit. Sequencing reads (both our new *G. sinensis* data and the previously-reported *S. mediterranea* data) were converted to fastq format using bcl2fastq2, quality controlled using FastQC, trimmed using TrimGalore, quality controlled again using FastQC and mapped to the *S. mediterranea* or *G. sinensis* transcriptome using Trinity script align_and_estimate_abundance.pl with aln_method=bowtie2 and est_method=RSEM.

The distribution of the log-transformed expression data is well represented by the superposition of two Gaussians (Supp. Fig. 4A-B). We associate the Gaussian with the lower mean with noise and the Gaussian with the higher mean with a biologically meaningful signal. This allows us to define a threshold (2 for *S. mediterranea* and 0.6 for *G. sinensis*, solid black lines). We only consider genes, for which the expression is above the threshold in 3 consecutive worm slices. Furthermore, we restrict our analysis to genes, which show a large expression change in comparison to the fluctuation between adjacent slices. For this, we compute the mean expression over a sliding window of 3 data points and determine the standard deviation relative to this smoothed curve. The difference between lowest and largest expression value generally increases for genes with increasing standard deviation (Supp. Fig. 4C-D). This allows us to define a threshold to exclude genes with small overall change relative to the fluctuations (we choose the difference between maximum and minimum value to be at least 5x larger than the standard deviation about the smoothed trend, solid black line). Next we perform complete hierarchical clustering based on the correlation distance. We determine a cut-off from the behaviour of cluster numbers (Supp. Fig. 4E-F). For small cut-off values, there are only very few non-trivial clusters, which combine only the most highly correlated genes. With increasing cut-off, there are more clusters and the clusters are also growing in size. Yet, there is a maximum, after which the cluster number decreases again and eventually all genes are combined into one giant cluster. We observe a small plateau / inflection point at 13 clusters for *S. mediterranea* and 17 clusters for *G. sinensis*, respectively (solid black lines), for which the cluster number is stable for a larger range in the cut-off value, before it rapidly decreases again. We assume that this provides a set of meaningful clusters, which we sort based on the position of the maximum expression value (Fig. 3D-E).

#### Identification and amplification of planarian orthologous genes

*G. sinensis* orthologues of *S. mediterranea* genes were identified as follows. First, a deposited *S. mediterranea* cDNA sequence corresponding to a gene of interest was downloaded from Genbank, imported into the Geneious software and queried against the reference *S. mediterranea* transcriptome assembly to identify the full-length cDNA sequence. Second, the longest open reading frame (ORF) was identified and translated to get an amino acid sequence that could be used as input for blastp. Third, the translation was queried against our new *G. sinensis* transcriptome assembly to identify potential *G. sinensis* orthologues ranked according to degree of pairwise identity. Fourth, the top hit was verified as a bona fide *G. sinensis* orthologue of the desired *S. mediterranea* gene by reciprocal BLAST to the *S. mediterranea* transcriptome. All orthologues are listed in Supp. Table 1.

Species-specific cDNA libraries were synthesised from total RNA using the SuperScript III First-Strand Synthesis System exactly according to manufacturer’s instructions. Target sequences were amplified from cDNA using Phusion DNA polymerase and primers listed in Supp. Table 2. Most amplified sequences were inserted by ligation-independent cloning^49^ into the pPRT4P^50^ vector. *S. mediterranea β-catenin-1* and *ef1a* were alternatively cloned into the paff8cT4P vector^51^ or pHAT2-T4P vector for protein expression. For certain genes, we bypassed cloning by adding universal overhangs during cDNA amplification, followed by a second round of PCR to introduce the T7 primer sites necessary for downstream in vitro transcription. In all cases, the desired target’s successful amplification was confirmed through Sanger DNA sequencing, either of the final plasmid when cloning was performed or the in vitro transcription template when cloning was not performed.

#### Antibody production

All steps of antibody production were carried out as previously described^16^. Recombinant fragments of S. mediterranea β-catenin-1 and EF1A were expressed in E. coli and purified under denaturing conditions. BALB/c mice were immunised with each antigen separately, and those with sera that detected the target planarian protein by Western blotting were sacrificed; their splenocytes were then used to create clonal hybridoma cell lines. Clone supernatants were initially screened using the Meso Scale Discovery platform, and promising candidates were further evaluated by Western blotting. Positive clones were expanded, and the antibodies were purified from the supernatant, acid-eluted, and buffer-exchanged into PBS. Clones 3588-D84-1 and 3621-D83-3 were identified as high-quality binders for anti-β-catenin-1 and anti-EF1A, respectively, based on the presence of an RNAi-sensitive band at the expected size and minimal background reactivity.

#### Quantitative Western Blotting

Animals were fixed and stored for several days to weeks at −80°C in a custom zinc-based fixative (100 mM ZnCl₂ in 100% ethanol)^52^, until ready for protein isolation. For analyses of β-catenin-1 abundance along the A/P axis, animals were transferred to a Peltier-cooled surface set at 4°C and then cut into six equal-length sections using a stab knife. The sections were pooled into six different 1.5 ml tubes according to their A/P position. Residual zinc fixative was removed from whole worms or tissue sections, and 120 µl of urea-based lysis buffer (9 M urea, 100 mM NaH₂PO₄, 10 mM Tris, 2% SDS, 130 mM DTT, 1 mM MgCl₂, pH 8.0) supplemented with 0.01 volume of Benzonase and 0.01 volume of protease inhibitor cocktail was added. Samples were left at room temperature for 20 minutes to allow the lysis buffer to fully permeate the tissue, then homogenised using a pellet mixer until no tissue was visible, and left at room temperature for an additional 20 minutes to ensure complete homogenization. For samples with a relatively large tissue volume, extra lysis buffer was added after mechanical homogenization to maintain an adequate lysis buffer-to-tissue ratio. Samples were then centrifuged at maximum speed (21,000 rcf) for 10 minutes at room temperature to pellet any insoluble matter, and supernatants were transferred to fresh tubes. A 2 µl aliquot from each sample was diluted 1:10 in autoclaved water and measured using a Nanodrop with the Protein A280 function, using a 1:10 dilution of the lysis buffer as a blank. Samples were then supplemented with homemade 5x lithium dodecyl sulfate (LDS; 530 mM Tris-HCl, 700 mM Tris Base, 10% w/v LDS, 50% w/v Glycerol, 2.55 mM EDTA, 500 mM DTT, 0.110 mM SERVA Blue G250, 0.875 mM Phenol Red, pH 8.5) sample buffer to reach a final concentration of 1x, followed by vortexing, aliquoting, and storage at −80°C.

Before loading onto the gel, samples were thawed, vortexed, heated to 70°C for 10 minutes, vortexed again, and spun down using a benchtop centrifuge. The samples were then run on 10-, 12-, 15-, or 17-well NuPage 4-12% Bis-Tris gels using XCell SureLock MiniCell devices set to a constant voltage of 120V and a homemade MOPS-based running buffer (50 mM MOPS, 50 mM Tris Base, 0.1% SDS, 1 mM EDTA, 5 mM NaHSO₃, pH 7.7). Following gel electrophoresis, the proteins were transferred onto a nitrocellulose membrane using the XCell II Blot module at a constant 30V with a pre-chilled, homemade NuPage transfer buffer (25 mM Bicine, 25 mM Bis-Tris, 1 mM EDTA, 0.05 mM chlorobutanol, 0.01% SDS, pH 7.2). When a large number of samples needed to be analysed on a single membrane (e.g., for gradient scaling experiments), the gels were cut with a scalpel after SDS-PAGE to isolate the regions where β-catenin-1 and EF1A bands were located. Four gel pieces containing β-catenin-1 or EF1A were then transferred together onto one membrane, allowing more samples to be compared on a single blot than would be possible without cutting (such as on a 17-well gel). After transfer, the membranes were dried at 37°C for 30 minutes and, if necessary, cut into sections for probing with different antibodies from the same species.

Membranes were blocked in blocking solution (5% w/v soy protein isolate in PBS) for 1 hour at room temperature with agitation. The detection procedure was carried out twice: first for β-catenin-1 and then for EF1A, since both primary antibodies were raised in mice. Membranes were incubated in β-catenin-1 primary antibody solution (0.2 µg/ml anti-β-catenin-1 in 0.5% soy protein isolate) overnight at 4°C with agitation, followed by three rinses with PBS and three washes for 10 minutes each with PBS + 0.1% Tween 20 (PBST). Membranes were then incubated in a secondary antibody solution (IRDye 800CW or CF770, diluted 1:20000 in 5% soy protein isolate) in the dark at room temperature with agitation. After three rinses with PBS, three 10-minute washes with PBST, and a final 15-minute wash with PBS, membranes were dried at 37°C for 20 minutes. Dried blots were imaged using an Odyssey Sa or Typhoon imager. Subsequently, membranes were rehydrated in PBS and incubated overnight in the EF1A primary antibody solution (0.03 µg/ml anti-EF1A in 0.5% soy protein isolate), and the remaining steps were repeated as done for β-catenin-1. Background-subtracted signal intensities were measured using Image Studio Lite or ImageQuant TL software, and the values were imported into an Excel spreadsheet for further analysis. The β-catenin-1 signal was normalised first to its respective EF1A signal to adjust for variations in protein loading, and then normalised again to one reference sample (as indicated in the figure legends) to facilitate data plotting.

#### RNA interference

The general strategy for inducing RNAi involved feeding planaria liver paste mixed with in vitro-synthesised double-stranded RNA (dsRNA) complementary to the target endogenous mRNA, largely as previously described^53^. dsDNA templates containing flanking T7 promoter sites, necessary for in vitro transcription, were generated by PCR from target sequences cloned into the pPRT4P vector. In vitro transcription was carried out in 200 µl reactions consisting of 8 µg of column-purified DNA template, 1x transcription buffer (0.4 M Tris pH 8.0, 0.1 M MgCl₂, 0.002 M spermidine, 0.1 M DTT), 1.5 mM Tris-buffered NTPs, 200 units of RiboLock RNase inhibitor, 0.3 units of inorganic pyrophosphatase, 170 units of T7 polymerase, and autoclaved water, incubated at 37°C overnight. The resulting dsRNA was annealed by heating at 95°C for 3 minutes and slowly cooling to room temperature using a ThermoShaker. It was then precipitated by mixing 1:1 with a precipitation solution (2.5 mM NaCl, 20% polyethylene glycol, 10 mM Tris-HCl pH 8.0) and centrifuged at 21,000 rcf at 4°C for more than 30 minutes. The pellet was washed twice with 70% ethanol, briefly air-dried, and resuspended in autoclaved water. The concentration of dsRNA was determined using a Nanodrop (with a custom conversion factor of 45), and its quality (verified by the presence of a single band at the expected size) was assessed by agarose gel electrophoresis. To prepare RNAi food, 12-16 µg/µl dsRNA solutions were mixed 1:4 with liver paste to achieve a final working concentration of 3-4 µg dsRNA/µl of total food volume. This mixture was aliquoted and stored at −80°C. RNAi was induced by feeding the planaria six times over a 16-day period, followed by a 7-day starvation period.

#### In situ hybridisation (ISH)

Sample preparation was carried out largely as previously detailed^54^ with several modifications. Intact and regenerating specimens were euthanized in 5% NAC (10% for fully-regenerated pieces from 18 mm *G. sinensis*) diluted in 1x PBS (supplemented with 0.1% MgCl₂ and 0.1% Tween-20 for *G. sinensis*), fixed in 4% PFA for 30 minutes (60 minutes for *G. sinensis*), and washed in 1x PBS with 0.3% (v/v) Triton-X-100 (PBSTx). Fully regenerated pieces from 18 mm *G. sinensis* were further treated with a “reduction solution” (1% NP-40, 1% SDS, 50 mM DTT, 1x PBS, brought to volume with autoclaved H₂O) at 37°C for 10 minutes. Samples were dehydrated in 50% (v/v) MeOH/PBSTx, followed by 100% MeOH, and stored at −20°C until FISH.

Riboprobe synthesis was also largely performed as previously described^54^. dsDNA templates containing a single flanking T7 promoter were generated by PCR from cloned or amplified planarian sequences. In vitro transcription was performed using 20 µl reactions containing >1 µg of column-purified DNA template, 1x transcription buffer, 1x DIG RNA labelling mix, 40 units of RiboLock RNase inhibitors, 0.03 units of inorganic pyrophosphatase, 40 units of T7 polymerase, and nuclease-free water, incubated at 37°C overnight. Following transcription, 1 µl of RNase-free DNase was added to remove template DNA by incubation at 37°C. Single-stranded RNA was precipitated by adding 0.75 volume of chilled 7.5 M ammonium acetate and 2 volumes of 100% ethanol, vortexed briefly, incubated at −80°C for over 30 minutes, and centrifuged at 21,000 rcf at 4°C for over 30 minutes. The samples were washed twice with 70% ethanol, air-dried briefly, and resuspended in 100 µl of deionized formamide.

The ISH proper was also largely performed as previously described^54^. Samples were transferred to 12-well plates, rehydrated in 50% MeOH/PBSTx, then PBSTx, and equilibrated in 1x saline-sodium citrate (SSC). Bleaching was done using a bleaching solution (5% non-deionized formamide, 1.2% H₂O₂, 0.5X SSC, autoclaved water to volume) on a light box for 2 hours, followed by PBSTx washes. Samples were treated with 2 µg/ml Proteinase K in 0.1% SDS for 10 minutes, post-fixed in 4% PFA for 10 minutes, washed in PBSTx, and equilibrated in “PreHyb” solution (50% formamide, 5X SSC, 1X Denhardt’s Solution, 100 µg/µl heparin, 1% Tween-20, 1 mg/ml torula yeast RNA, 50 mM DTT) for 2 hours at 56°C. Hybridization was carried out by incubating samples overnight (>12 hours) at 56°C in “riboprobe mix” (riboprobes diluted 1:3000-1:5000 in “Hyb” solution: 50% formamide, 5X SSC, 1X Denhardt’s Solution, 100 µg/µl heparin, 1% Tween-20, 0.25 mg/ml torula yeast RNA, 50 mM DTT, 5% w/v dextran sulfate) within a humidified hybridization oven. To prevent evaporation-associated artifacts, water was added to empty wells and the space between wells and the plate was double-sealed prior to overnight incubation. Post-hybridization, samples were washed twice for 20 minutes with “Wash Hyb” (50% formamide, 0.5% Tween-20, 5X SSC, 1x Denhardt’s), twice for 20 minutes with 0.5X Wash Hyb + 1X SSC, three times for 20 minutes with 2x SSC, and three times for 20 minutes with 0.2x SSC, all at 56°C. After cooling to room temperature, samples were washed three times for 5 minutes with PBSTx and incubated in a blocking solution (5% sterile horse serum + 0.5% Roche Western Blocking Reagent in PBSTx) for 1 hour at room temperature. Next, samples were incubated overnight at 4°C in an antibody solution (1:2000 anti-DIG-POD diluted in blocking solution).

For fluorescent detection (all targets except *G. sinensis sfrp1*), samples were washed 6x 20 min with PBSTx, then incubated for 30 minutes at room temperature in TSA buffer (2M NaCl, 0.1M Boric acid pH 8.5) with 0.006% H₂O₂, 1:3000 Sulfo-Cyanine 3 tyramide, and 20 µg/ml 4-IPBA peroxidase enhancer (in the dark). From this point on, samples were exposed to minimal light to preserve fluorescent signals. They were washed in PBSTx twice for 5 minutes at room temperature, then incubated overnight at 4°C in 1:2000 DAPI diluted in PBSTx, post-fixed for 20 minutes in 4% PFA/PBSTx at room temperature, washed twice for 5 minutes, and cleared in Scale S4 (10% glycerol, 15% DMSO, 40% sorbitol, 4M urea, 0.1% Triton X-100)^55^. For colorimetric detection (*G. sinensis sfrp1*), samples were washed 6x 20 min with TNTx, then incubated in “AP” solution (0.1M Tris pH 9.5, 0.1M NaCl, 0.05M MgCl_2_, 1% Tween-20) for 10 min, AP supplemented with 0.27% NBT (Roche) and 0.53% BCIP (Roche) substrates for 10 min and “DEV” solution (0.1M Tris pH 9.5, 0.1M NaCl, 0.05M MgCl_2_, 1% Tween-20, 7.8% PVA, 0.27% NBT, 0.53% BCIP) for at least 30 min in the dark, until the expected expression pattern was unambiguous. Samples were washed in PBSTx 2x 5 min at RT then overnight at 4°C, post-fixed for 45 min in 4% PFA/PBSTx at RT, washed 2x 5 min at RT and cleared in “Scale A2” (2M Urea, 75% glycerol)^56^. Samples were mounted between two coverslips separated by a spacer made from a double-sided adhesive sheet to prevent squashing and stored at 4°C until imaging.

#### Light microscopy

Images of intact and regenerating living animals were captured using a custom-made macrophotography setup, which included an EOS 6D camera fitted with a macro objective lens and mounted on a stand positioned above a light box. *G. sinensis wnt11-2* ISH images were taken at 10x magnification using a VS200 Research Slide Scanner. *G. sinensis sfrp1* ISH images were taken using a Nikon AZ100M microscope equipped with a Digital Sight DS-Fi1 camera. All other ISH images were obtained at 20x magnification with an IXplore Spin Confocal Imaging Microscope System. The acquisition settings were maintained consistently throughout each experiment.

#### Image processing and analysis

Brightfield images were processed using Adobe Photoshop CS5.1, including rotation, cropping, and brightness adjustments. Fluorescence image processing and analysis were conducted with Fiji. For the *notum* ISH images (Fig. 2), maximum intensity projections were created, and the images were rotated so that the anterior wound edge (determined by the orientation of the pharynx) faced left, then tightly cropped. The spatial distribution of *notum* expression along the A/P axis of each piece was quantified at sub-micron resolution using the “Plot Profile” function and divided into 10 equal bins. The degree of A/P symmetry was calculated by dividing the mean intensity of the anterior 5 bins by the mean intensity of the posterior 5 bins.

#### Pharmacology

Commercially available C59 was diluted to 10 mM in DMSO and stored in single-use aliquots at −80°C. Experimental drug solutions were prepared by mixing C59 stock (to achieve final concentrations of 0.4, 2, 10, or 50 µM as indicated in the figure legends) with supplemental DMSO (to a final concentration of 0.5% in *G. sinensis* experiments and 0.75% in *S. mediterranea* experiments) in planarian water, followed by vortexing at maximum speed for 30 seconds. For cWnt gradient perturbations, uninjured animals were maintained in the drug solution for 7-14 days (as specified in the figure legends), with daily exchanges of the solution. After the treatment period, the animals were transferred to a plastic container holding 1.1 litres of planarian water, which was continuously agitated by an aquarium air pump to facilitate the removal of C59 from the tissue. For post-amputation cWnt perturbations, freshly cut pieces were exposed to the drug solution for 48 hours (with a solution exchange after 24 hours) before being moved to fresh petri dishes with planarian water to wash out C59 from the tissue. In all experiments, vehicle-only control cohorts were maintained in planarian water supplemented with 0.5% (*G. sinensis*) or 0.75% (*S. mediterranea*) DMSO and otherwise treated identically to C59-treated ones.

#### Mass spectrometry

Pools of intact or regenerating *S. mediterranea* or *G. sinensis* were transferred to pre-weighed 1.5 ml Eppendorf tubes, rinsed once with planarian water, aspirated to remove excess liquid, and stored at −80°C. Samples were then dried for 1 hour at 37°C in a vacuum concentrator and reweighed. Homogenization involved adding 300 µl of mass spectrometry-grade MeOH containing 2 µM LGK-974 (used as an internal control to account for technical variability) and three 2.3 mm stainless steel beads to each sample. The samples were mechanically disrupted for 5 minutes at 30 Hz using a TissueLyser II. After homogenization, samples were centrifuged at 21,000 rcf and room temperature for 5 minutes, and the supernatants were purified using MicroSpin C18 columns according to the manufacturer’s instructions. Samples, hereafter referred to as “worm extracts”, were stored at −20°C until mass spectrometry analysis.

Immediately prior to mass spectrometric analysis, 5 µl of each worm extract was diluted 1:10 with spray mix (7.5 mM ammonium formate in a 4:2:1 v/v/v mixture of isopropanol:MeOH:chloroform) in wells of a skirted twin.tec PCR plate, in triplicate. To enable absolute quantification, five “standards”, generated by serially diluting pure C59 in untreated worm extract to a final concentration of 2560/5120/10240/20480/40960 nM, were also diluted 1:10 with spray mix and included on the plate in triplicate. Samples were introduced into a Q Exactive hybrid quadrupole Orbitrap mass spectrometer via TriVersa NanoMate automated ion source equipped with microfluidics ESI Chip. Mass spectrometric analysis was performed in positive mode under the following instrumental settings: running time 2.5 min/sample, capillary temperature 200°C, resolution *R_m_*_/*z*=200_ 140000, scan range m/z 200-500; lock mass 445.12003 and 338.34174. To confirm the identity of C59 and LGK-974 signals in worm extracts, MS2 was performed for the corresponding precursors with m/z 397.177 and 380.176 using mass isolation window of 3Da and normalised collision energy (NCE) 25 and 45 respectively and the spectra compared with fragmentation pattern of the commercial C59 standard. Raw data were first processed with PeakStrainer software^57^ to remove noise signals. To obtain the signal intensities for precursors of analytes of interest, C59 and LGK-974, the resulting files in *.mzXML format were processed by LipidXplorer software^58^.

Relative C59 tissue abundance was calculated as follows. First, C59 signal intensity was normalised to the LGK-974 signal intensity to correct for technical variability. Second, linear regression analysis was applied to the LGK-normalised C59 signals of the five standard samples to create an equation for determining the unknown absolute amount of C59 in each experimental sample. Third, the concentration of C59 in the tissue was calculated by dividing the total C59 amount by the dry weight of the sample, which was determined by subtracting the initial weight of the empty tube from the weight of the tube containing the dried tissue sample. Fourth, the concentration of each sample was normalised to a single sample from within the same experiment (the 0 days washout sample in Fig. 4I-J and the Pre 1 dpa sample in Supp. Fig. 3C) and converted to % to facilitate graphical representation.

#### Statistics

All statistical analyses were conducted using GraphPad Prism 10.0 (GraphPad Software, Inc.). The specific tests employed are detailed in the figure legends.

## Author contributions

JPC, HT-KV and JCR conceptualised the study, designed experiments and interpreted the results. JPC performed most experiments. HT-KV performed the β-catenin-1 measurements presented in Fig. 6. JEMD, ASc and ASh contributed to establishment of the pharmacological approach presented in Fig. 4. AR assembled the G. sinensis transcriptome and processed the RNA-seq data presented in Fig. 3. SW performed the RNA-seq analyses presented in Fig. 3. JPC and JCR wrote the manuscript with input from all authors. JCR supervised the study.

## Acknowledgements

We thank past and present members of the Rink lab for discussions and sharing of expertise, including Miquel Vila-Farré (planarian biodiversity), Tobias Boothe (microscopy), Markus Grohme (molecular biology), Albert Thommen (EF1A antibody establishment), Heino Andreas and Claudia Koch (wormcare support) and Stephanie von Kannen, Klaske Schippers and Delia Niehaus (lab management); Elly Tanaka, Pavel Tomancak and the late Suzanne Eaton for inspiring project discussions; MPI-CBG Antibody and Protein Biochemistry core facilities; and the DRESDEN-concept Genome Center for support with next-generation sequencing. JPC acknowledges funding from the Boehringer Ingelheim Fonds PhD fellowship program. JCR received funding from the European Research Council (ERC) under the European Union’s Horizon 2020 research and innovation program (grant agreement number 649024), from the German Research Foundation (project RI 2449/51), from the Behrens-Weise Foundation and from the Max Planck Society. Open access funding provided by Max Planck Society.

## Declaration of interests

The authors declare no competing interests.

## Supplemental material

**Supplemental Figure 1.**
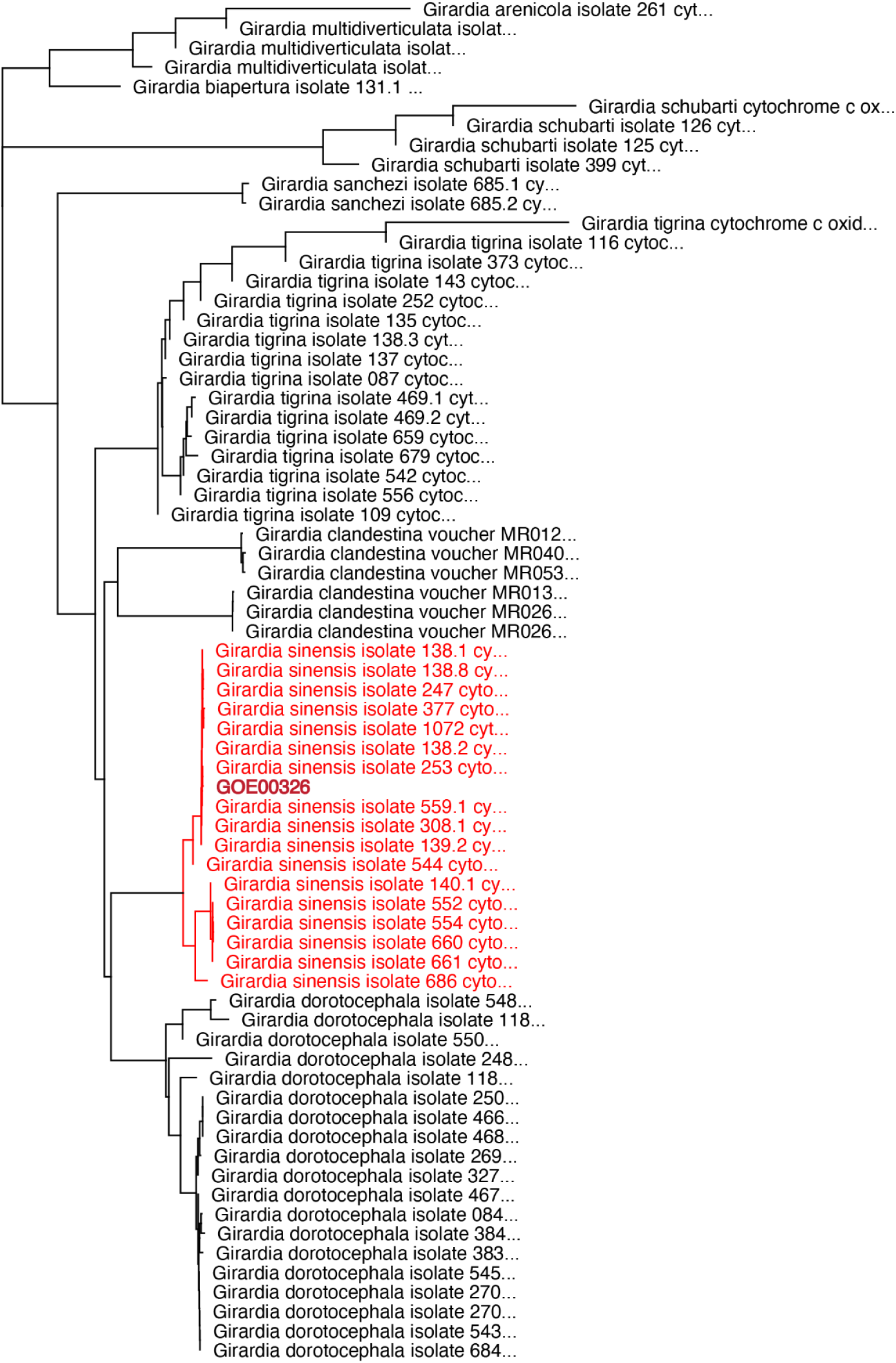
A distance-based phylogenetic tree of *cox1* sequences from nine different *Girardia* species^27^ and the *Girardia* strain used in this study, GOE00326. The placement of the GOE00326 *cox1* sequence is strong support for a *G. sinensis* species-level classification. See Materials and Methods for a detailed explanation of how the tree was generated.

**Supplemental Figure 2.**
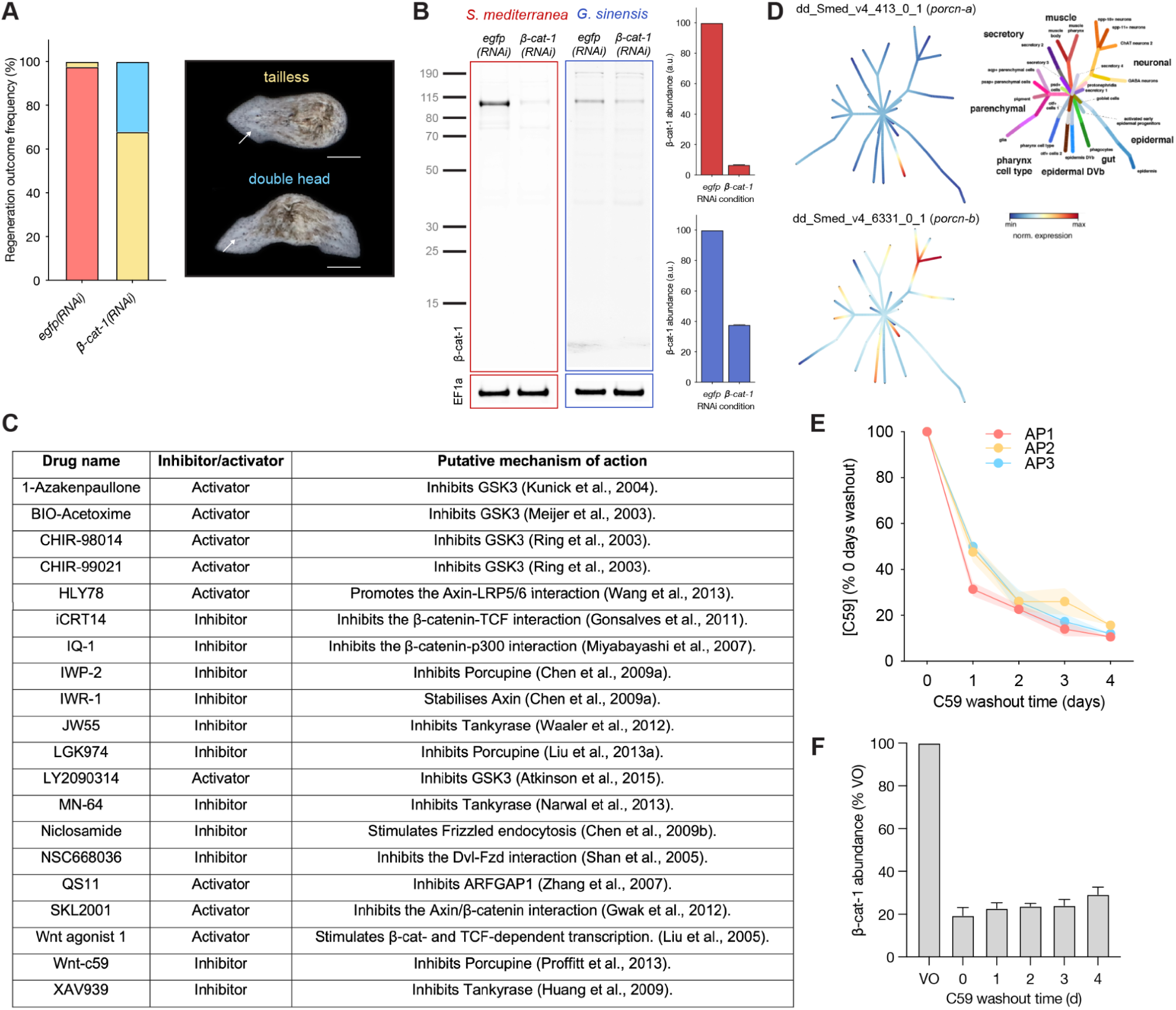
A) Quantification of regeneration outcomes in 1 mm pieces from 6 mm *S. mediterranea* after *β-catenin-1(RNAi)* or control *egfp(RNAi)*. N = 68 pieces from one representative experiment. White arrows indicate eye spots as morphological head marker. Scale bars: 500 µm. B) Quantitative Western blot measurements on *egfp(RNAi)* control and *β-catenin-1(RNAi)* protein extracts to verify specificity of the anti-β-catenin-1 antibody. EF1A was used as loading control. n = five individuals per RNAi condition. N = one representative experiment. C) List of screened commercially-available cWnt signalling inhibitors and activators. D) Graphical representation of the expression of two *S. mediterranea porcupine* homologues across planarian cell types. Generated with the Planaria Single Cell Atlas online resource (https://shiny.mdc-berlin.de/psca/)^34^. E) Clearance of C59 from the tissue of 18 mm *G. sinensis* treated for 7 days with 2 µM C59 and then subjected to 4 days of drug washout. AP1, AP2 and AP3 represent different positions along the A/P axis. n = 3 individuals per time point. N = three independent experiments. Error bands represent SEM. F) Quantitative Western blotting of cWnt signalling activity in the tail tip of 18 mm *G. sinensis* treated for 7 days with 2 µM C59 and then subjected to 0, 1, 2, 3 or 4 days of drug washout. Vehicle-only (VO) controls were treated for 7 days with DMSO and then subjected to 4 days of washout. Error bars represent SEM of three independent experiments. Each experiment is normalised to its respective VO measurement.

**Supplemental Figure 3.**
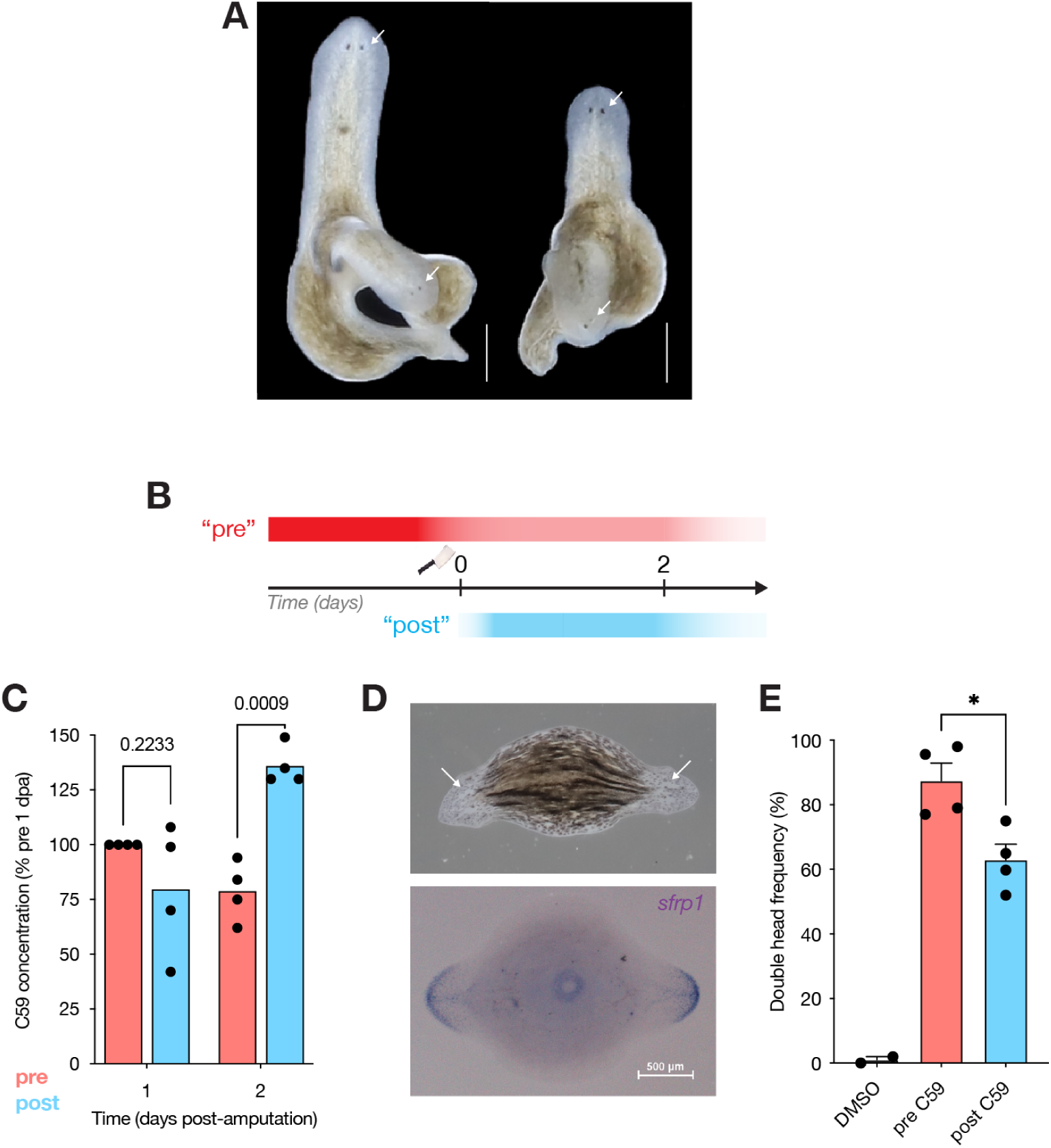
A) Representative live images of the two double-headed individuals resulting from the experiment shown in Fig. 5G-J. White arrows indicate eyes and thus head identity. Scale bars: 500 µm. B) Overview of experimental approach. “Pre” large *G. sinensis* were treated with 2 µM C59 or 0.5 % DMSO as vehicle-only control for 7d, washed for 1d to allow for partial drug clearance, cut into 9ths and left to regenerate. “Post” animals were untreated prior and instead treated for 2 d with 0.5 µM C59. The shaded red rectangle represents tissue C59 levels contributed by the gradient manipulation (dark red) and persisting from 0 days post-amputation (dpa) onwards (light red). The shaded blue rectangle signifies tissue C59 levels contributed exclusively by post-amputation treatment. C) Quantification of tissue C59 concentration in Pre vs Post regenerating pieces. Each point represents the average of three technical triplicate measurements, normalised to the respective Pre 1 dpa value. n = 14 pieces per data point. N = 4 independent experiments. D) Representative images of live morphology and expression pattern of the anterior marker *sfrp1* at 10 dpa. White arrows indicate eyes. In these experiments, regenerating pieces with biaxial expression of *sfrp1* were scored as double-headed.Quantification of double-head frequency. n = 369 Pre and 326 Post pieces. N = 4 independent experiments. Statistical significance was assessed by Wilcoxon test (D) or Mann-Whitney test (E). ns = not significant (i.e. p > 0.05), * = significant (i.e. p < 0.05).

**Supplemental Figure 4.**
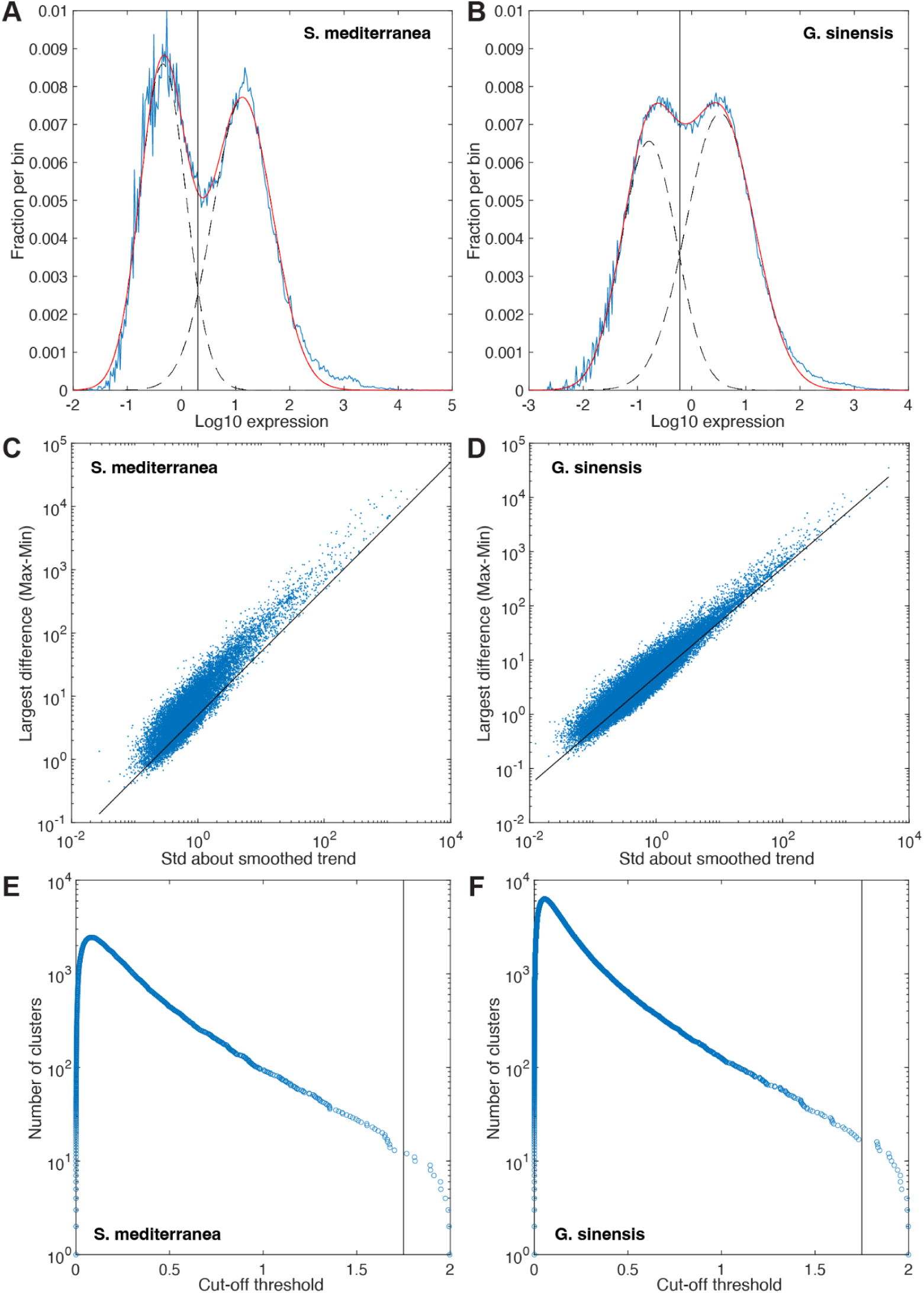
A-B) Log expression of RNA sequencing counts of *S. mediterranea* (left) and *G. sinensis* (right) shows two peaks, which are fitted by a linear combination (red) of two Gaussian functions (black dashed). We associate the Gaussian with the lower mean with noise, which allows us to define a threshold (black solid) to distinguish a meaningful signal from noise C-D) The difference between maximum and minimum expression of each gene (blue dots) is compared to the fluctuation (Std) about the smoothed trend (moving average of 3 data points). This allows us to define a threshold (black line) to identify genes, for which the maximum expression change is not much larger (<5x) than the average fluctuations about the smoothed trend E-F) Number of non-trivial clusters (i.e. with more than a single gene) for complete hierarchical clustering (with a correlation distance) is shown as a function of the distance cut-off. We define a threshold for our clustering (black solid) at an inflection of the curve, at which the cluster number is also stable for a range of cut-off values (i.e. gap between data points).

**Supplemental Video 1**

Representative double-head and tailless regeneration phenotypes observed in 2 mm long pieces from 18 mm long *G. sinensis* treated for 7 days with 2 µM C59, subjected to 4 days of drug washout, amputated and left to regenerate for 21 days. Related to Fig. 5O-R.

## Notes

### Competing Interest Statement

The authors have declared no competing interest.

### Summary of Updates

The statement "Open access funding provided by Max Planck Society" has been added to the acknowledgements section.

